# Differential abilities to engage inaccessible chromatin diversify vertebrate HOX binding patterns

**DOI:** 10.1101/2019.12.29.890335

**Authors:** Milica Bulajić, Divyanshi Srivastava, Jeremy S Dasen, Hynek Wichterle, Shaun Mahony, Esteban O Mazzoni

## Abstract

While *Hox* genes encode for conserved transcription factors (TFs), they are further divided into anterior, central, and posterior groups based on their DNA-binding domain similarity. The posterior *Hox* group expanded in the deuterostome clade and patterns caudal and distal structures. We aim to address how similar HOX TFs diverge to induce different positional identities. We studied HOX TF DNA-binding and regulatory activity during an *in vitro* motor neuron differentiation system that recapitulates embryonic development. We find diversity in the genomic binding profiles of different HOX TFs, even among the posterior group paralogs that share similar DNA binding domains. These differences in genomic binding are explained by differing abilities to bind to previously inaccessible sites. For example, the posterior group HOXC9 has a greater ability to bind occluded sites than the posterior HOXC10, producing different binding patterns and driving differential gene expression programs. From these results, we propose that the differential abilities of posterior HOX TFs to bind to previously inaccessible chromatin drive patterning diversification.

## INTRODUCTION

*Hox* genes encode a highly conserved transcription factor family that endows cells with positional identity during embryonic development (Duboule & Dolle, 1989; Lewis, 1978; McGinnis & Krumlauf, 1992). In mammals, *Hox* genes are organized into four clusters located on different chromosomes (*HoxA, HoxB, HoxC*, and *HoxD)*. Each cluster contains a subset of 13 similar paralogous *Hox* genes, genomically arranged in the same linear order as their spatial and temporal expression patterns in the developing embryo, a phenomenon known as collinearity (Duboule & Morata, 1994; Kmita & Duboule, 2003). Changes in *Hox* gene expression patterns induce gross morphological changes, resulting in well-characterized homeotic transformations. However, how HOX transcription factors (TFs) assign different positional identities during cell differentiation is not entirely understood.

*Hox* genes encode for similar homeodomain-containing TFs (Akam, 1989; Regulski et al., 1985). Homeodomains are highly conserved helix turn helix DNA-binding domains that recognize similar consensus DNA sequences (Affolter, Slattery, & Mann, 2008; Berger et al., 2008; Gehring et al., 1994; Noyes et al., 2008). Vertebrate HOX TFs can be divided into anterior (HOX1-5), central (HOX6-8), and posterior (HOX9-13) paralog groups. The posterior HOX group is particularly interesting because it has expanded in the deuterostome clade. While there is one *Drosophila Abd-B*, several *Abd-B*-related *Hox* genes assign different posterior positional identities during vertebrate development (reviewed in (Duboule, 2007; Lanfear, 2010)). Thus, understanding how paralogous HOX TFs differentiate their genomic binding activity to specify cell fates is at the core of understanding vertebrate body patterning.

*In vitro* binding studies have investigated the intrinsic sequence preferences of HOX TFs alone or in complex with specific cofactors. These studies demonstrate that the anterior (HOX1-5) and central HOX paralog groups (HOX6-8) prefer to bind motifs containing the canonical TAAT core sequence, whereas the posterior paralog groups (HOX9-13) preferentially bind TTAT core sequences (Berger et al., 2008; Ekker et al., 1994; Mann, Lelli, & Joshi, 2009; Noyes et al., 2008). Moreover, the interaction between HOX TFs and MEIS and PBX cofactors increases the specificity and selectivity of HOX DNA-binding (reviewed in (Mann & Affolter, 1998; Mann & Chan, 1996; Merabet & Mann, 2016)). However, how vertebrate HOX TFs within a single group diversify their genomic binding patterns remains obscure.

Despite the extensive analysis of HOX TF binding *in vitro*, relatively little is known about HOX binding specificity in the context of cellular chromatin landscapes (De Kumar et al., 2017; Donaldson et al., 2012; Huang et al., 2012). For example, virally-expressed HOXA9-13 and HOXD9-13 in primary chicken mesenchymal limb progenitors exhibit some binding specificity differences between the posterior group HOX TFs, with HOXA/D13 paralogs being the most different (Jerkovic et al., 2017). In *Drosophila*, recent studies have investigated the role of chromatin accessibility in shaping Hox TF binding. Of the central and posterior fly Hox factors, *Drosophila* Abd-B displays an increased ability to bind previously inaccessible chromatin (Beh et al., 2016; Porcelli, Fischer, Russell, & White, 2019). However, it is not possible to investigate how binding selectivity has diverged between the vertebrate Abd-B-derived posterior HOX TFs (HOX9-HOX13) using *Drosophila* models. In line with their differential patterning activities, the vertebrate HOX TFs may have diverged either in their sequence preferences or in their differential abilities to engage inaccessible chromatin.

In vertebrates, *Hox* genes pattern various developing tissues. Notably, spinal cord neuronal diversity requires *Hox* gene activity along its rostro-caudal axis (Dasen & Jessell, 2009; Dasen, Liu, & Jessell, 2003; Sweeney et al., 2018). The role of *Hox* genes in organizing the spinal cord presents an interesting conundrum. The limb-innervating expression program is controlled by central (HOX6 & HOX8) HOX TFs at the brachial level and posterior (HOX10) HOX TFs at the lumbar spinal cord (Dasen, De Camilli, Wang, Tucker, & Jessell, 2008; Jung et al., 2018; Lacombe et al., 2013; Rousso, Gaber, Wellik, Morrisey, & Novitch, 2008; Wu, Wang, Scott, & Capecchi, 2008). Thus, a similar motor neuron fate is induced by HOX TFs with different DNA sequence preferences. Meanwhile, the posterior HOXC9 induces thoracic fate (Jung et al., 2010; Jung et al., 2014).

Thus, two posterior group genes, *Hoxc9* and *Hoxc10*, induce different spinal cord fates. In agreement with their genomic cluster position, *Hox13* paralogs are expressed late during development, distally, and in posterior regions. Moreover, they are associated with the unique ability to serve as “repressors of elongation” or “patterning terminators” by inhibiting cell growth or inducing apoptosis (Denans, Iimura, & Pourquie, 2015; Economides, Zeltser, & Capecchi, 2003; Godwin & Capecchi, 1998; Young et al., 2009). As a model to understand the differential patterning activities of HOX TFs, we sought to understand how central and posterior HOX TFs bind the genome to induce different spinal cord identities.

The study of genome-wide HOX TF binding in cellular contexts is challenging due to the lack of availability of homogenous relevant cell populations at scales compatible with chromatin immunoprecipitation (ChIP), and due to the limited ability to conduct temporal TF binding during development. To mitigate this technical limitation, we opted for an embryonic stem cell (ESC) differentiation system that recapitulates ventral spinal cord early development (Davis-Dusenbery, Williams, Klim, & Eggan, 2014; Wichterle, Lieberam, Porter, & Jessell, 2002; Wichterle & Peljto, 2008). In response to the dorso-ventral Hedgehog and rostro-caudal retinoic acid (RA) patterning signals, ESCs differentiate into motor neurons (MNs) and interneurons by transitioning through embryonic progenitor states (Wichterle et al., 2002). The culture acquires a rostral spinal cord identity, with 90% of cells expressing *Hoxa5* (Peljto, Dasen, Mazzoni, Jessell, & Wichterle, 2010; Peljto & Wichterle, 2011). Moreover, differentiating ESC-derived MNs respond to *Hox* gene overexpression similarly to those in the developing spinal cord (Jung et al., 2010; Machado et al., 2014; Narendra, Bulajic, Dekker, Mazzoni, & Reinberg, 2016; Tan, Mazzoni, & Wichterle, 2016). Thus, ESC-to-MN differentiation recapitulates critical aspects of MN differentiation and constitutes a suitable model to study HOX TF activity in relevant cellular and chromatin environments.

To understand how HOX TFs control different fates, we induced individual HOX TFs in progenitor motor neurons. We analyzed the differential binding of the HOX TFs in the context of their underlying sequence motifs and interactions with the preexisting chromatin accessibility environment. We focused on a subset of HOXC TFs (HOXC6, 8, 9, and 10) due to their importance in inducing spinal cord identities. We complemented these studies by examining additional HOX9 paralogs and the posterior HOXC13. Our results suggest that limb-innervating fate is not the product of identical central and posterior HOX binding patterns since HOXC10 does not mimic HOXC6 and HOXC8 binding profiles. Although posterior group HOX TFs have similar DNA motif preferences, they do not bind to the genome with identical patterns. This difference is mainly due to their differing abilities to bind to motifs occluded in inaccessible chromatin. For example, while HOXC9 and HOXC10 bind similar DNA sequence motifs, HOXC9 has a greater ability to bind previously inaccessible chromatin and thus accesses different binding sites. In summary, our work describes divergence in the abilities to engage inaccessible chromatin among vertebrate posterior group HOX factors derived from a single *Drosophila* gene. From these results, we propose that the differential abilities of posterior HOX TFs to bind to previously inaccessible chromatin is the predominant force driving their patterning diversification.

## RESULTS

### HOX TF expression controls neuronal fates during *in vitro* spinal cord differentiation

HOX proteins have similar DNA binding domains, yet they control positional identity along the rostro-caudal axis. In particular, posterior HOXC9 and HOXC10 have a single shared *Drosophila* ortholog, Abd-B, yet they pattern different spinal cord fates. We focus our attention on a subset of *HoxC* genes due to their cardinal spinal cord patterning activities (Dasen & Jessell, 2009; Dasen et al., 2003; Jung et al., 2010; Wu et al., 2008). We generated isogenic ESC lines that express *Hoxc6* (brachial), *Hoxc8* (brachial), *Hoxc9* (thoracic) or *Hoxc10* (lumbar) upon Doxycycline (Dox) treatment (Figure 1A, B, Supp Figure 1A) (Iacovino et al., 2011; Mazzoni et al., 2011). To recapitulate *Hox* expression dynamics during development, Dox was added at a time point when the culture consists of neural progenitors (Figure 1A, Supp Figure 1A).

**Figure 1:**
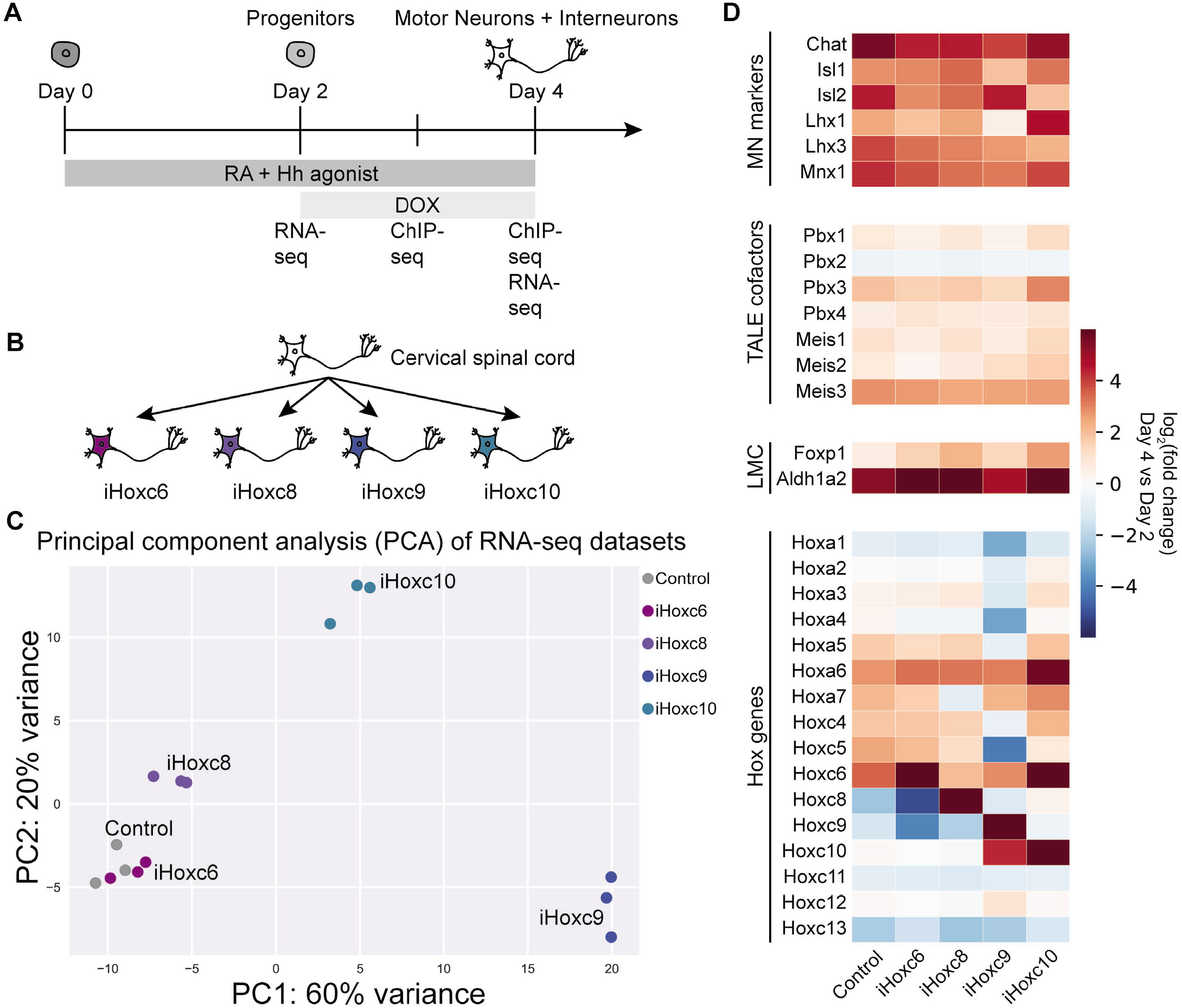
HOX TFs control cell fates during *in vitro* spinal cord differentiation. (A) Experimental scheme. (B) Diagram of neuronal cell fate control by *Hox* gene overexpression. (C) Principal Component Analysis (PCA) of the RNA-seq datasets (Day 4) reveals similarities in the gene expression profiles induced by HOX TFs. (D) RNA–seq heatmap showing the expression of representative marker genes in No Dox control, iHoxc6, iHoxc8, iHoxc9 and iHoxc10 neurons relative to Day 2 progenitors.

As time progresses, cells become neurons, peaking at 48h after Dox addition when the culture mostly consists of post-mitotic MNs as well as interneurons. We refer to these as iHoxc6, iHoxc8, iHoxc9, and iHoxc10 neurons, respectively. Importantly, all HOX proteins are N-terminally Flag-tagged, which allows for immunoprecipitation with the same antibody, eliminating any bias that could occur from different antibody affinities and cross-reactivity.

HOXC6, HOXC9, and HOXC10 divide the spinal cord into three important levels: brachial, thoracic, and lumbar, respectively. *Hoxc6*, *Hoxc8*, and *Hoxc9* inductions during *in vitro* motor neuron differentiation have been individually characterized with expression changes in a few downstream genes, and they produce the expected phenotypes (Jung et al., 2010; Narendra et al., 2016; Tan et al., 2016). To have a comprehensive and comparative integration of HOX expression consequences, we performed RNA-seq in *Hoxc6, Hoxc8, Hoxc9, Hoxc10* induced post-mitotic neurons as well as control neurons not treated with Dox (no Dox control) (Figure 1A, Supp Figure 1A). In a principal component analysis (PCA), the first two principal components explained 80% of the variance in RNA-seq tag counts, reflecting a combination of the paralog group and the subtype identities specified by the HOX proteins (Figure 1C) (see Methods). Control cells express *Hox* genes up to paralogs *Hox5*. Thus, iHoxc6 induces some expression changes grouping close to control (Figure 1C). The slightly more posterior inducing *Hox*, iHoxc8, separates along PC2. iHoxc9 grouped the furthest away, while iHoxc10 grouped between cells expressing *Hox5-8* genes (control cells, iHoxc6 and iHoxc8) and iHoxc9. Multidimensional scaling (MDS), a non-linear dimensionality reduction technique, produced a similar lower-dimensional embedding separating well between each inducible *Hox* line (Supp Figure 1B). Overall, this data show that HOX TFs induce distinct gene expression profiles during *in vitro* spinal cord differentiation, with an increasing transcriptional diversity by posterior HOX TFs (Figure 1C, Supp Figure 1B, C).

Although there is no global gene expression characterization of *Hox* expression manipulation during embryonic development, iHox lines induced the expression of known marker genes in agreement with previous studies. For example, *Hoxc6, Hoxc8*, and *Hoxc10* overexpression induce canonical LMC markers *Raldh2* and *FoxP1* (Figure 1D, Supp Figure 2A, B) (Narendra et al., 2016; Tan et al., 2016). *Hoxc9* overexpression leads to the repression of anterior *Hox1-5* paralogs (Figure 1D, Supp Figure 2A, B) (Jung et al., 2010). The various induced *Hox* genes display similar RNA levels in our RNA-seq experiments (represented as FPKM values), and these levels are also comparable to those of endogenous *Hoxc5* expressed in the control (No Dox control) (Supp Figure 2C). Importantly, *Hox* induction during *in vitro* spinal cord differentiation did not derail the ability of cells to acquire an MN identity (Figure 1D, Supp Figure 2A, B). Thus, HOX TF activity during ESC differentiation induces distinct spinal cord fates, recapitulating aspects of embryonic differentiation.

### HOXC6, HOXC9, and HOXC10 TFs have different genome-wide binding profiles

To understand how HOX TFs assign positional identity, we assayed the genomic binding of HOXC6, HOXC8, HOXC9, and HOXC10 by performing ChIP-seq experiments 24h and 48h after Dox treatment, newborn and young post-mitotic MNs, respectively (Figure 1A, Supp Figure 1A). These time points are critical for HOX positional identity patterning because HOX TFs control motor neuron types at early postmitotic states (Dasen et al., 2003). We restricted the analysis to the top 10,000 ChIP-seq sites in each dataset for all downstream analyses to ensure the least amount of bias from comparing different experiments and performed differential binding analysis via MultiGPS (Mahony et al., 2014) (which runs edgeR (Robinson, McCarthy, & Smyth, 2010) internally).

Despite sharing 82% similarity within their DNA-binding homeodomains (Supp Figure 3A), we find that the posterior HOXC9 and HOXC10 display divergent genomic binding patterns (Figure 2A, Supp Figure 3B). HOXC10 primarily binds a subset of HOXC9 binding sites; while 90% of HOXC10’s top 10,000 binding sites show similar enrichment levels for HOXC9 and HOXC10, an additional 5,230 sites are bound preferentially by HOXC9 (Figure 2A) (edgeR q-value <0.01, see Methods). Thus, although HOXC9 and HOXC10 contain similar DNA-binding domains, HOXC9 binds to additional genomic locations.

**Figure 2:**
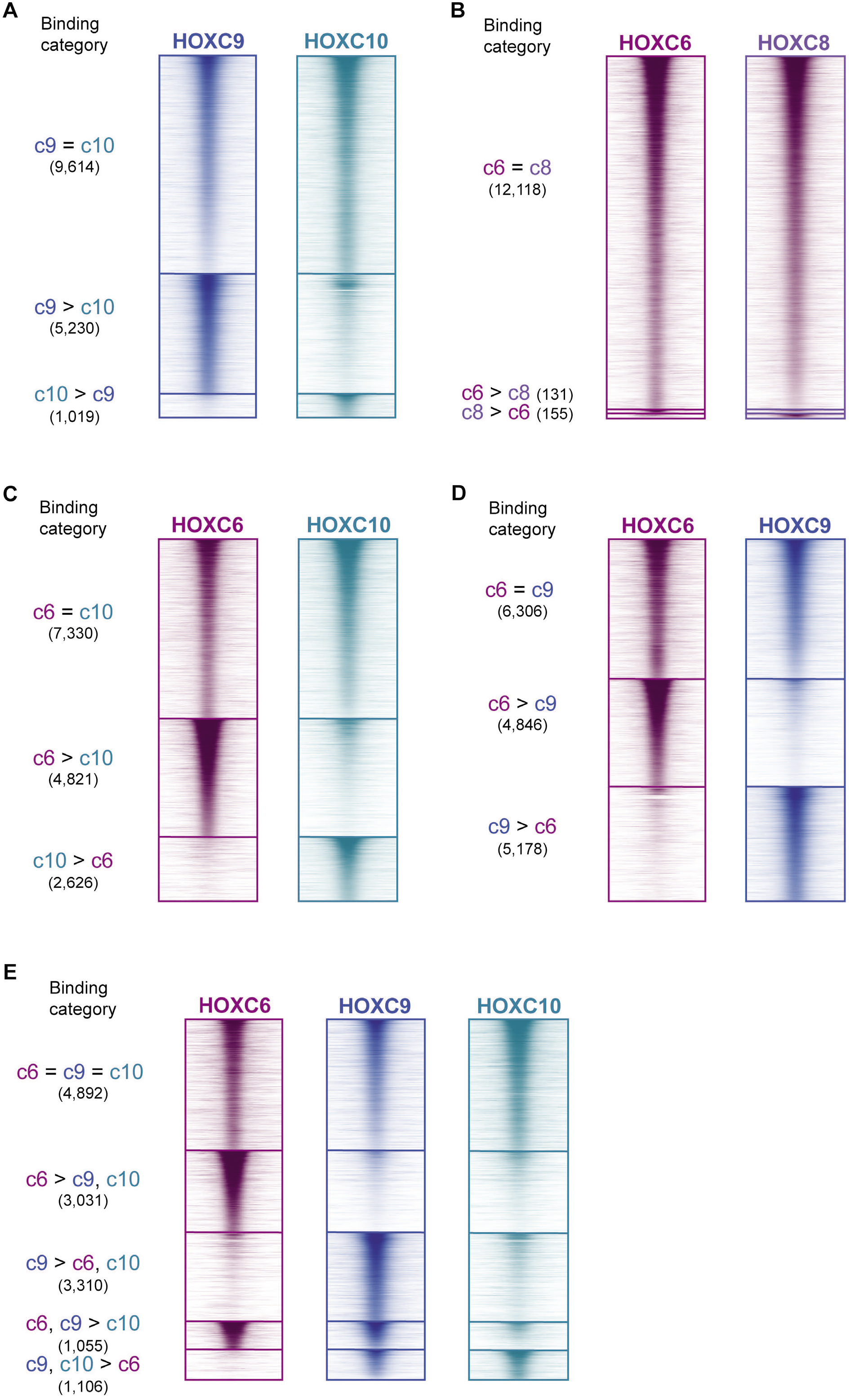
HOXC6, HOXC9, and HOXC10 TFs have different genome-wide binding profiles. (A-D) ChIP-seq heatmap showing binding comparisons of HOX TFs in differentiating neurons, at Day 3. Sites bound by both indicated HOX TFs noted as “=” sites. Preferentially bound sites by HOXC6, HOXC8, HOXC9 or HOXC10 noted as “c6 >”, “c8 >”, “c9 >”, and “c10 >”, respectively. (E) ChIP-seq heatmap showing binding comparisons of HOXC6, HOXC9, and HOXC10 in differentiating neurons, at Day 3. Sites bound by all three HOX TFs noted as “c6 = c9 = c10” sites. Preferentially bound sites by HOXC6, HOXC9, “HOXC6 and HOXC9”, or “HOXC9 and HOXC10” noted as “c6 > c9, c10”, “c9 > c6, c10”, “c6, c9 > c10”, and “c9, c10 > c6”, respectively.

We wondered if another pair of similar HOX TFs would also diverge in binding patterns. Thus, we compared the binding of central HOXC6 and HOXC8 TFs. HOXC6 and HOXC8 display few differences in enrichment at their top-most bound sites; fewer than 2% of tested HOXC6 and HOXC8 sites are significantly differentially bound by the two TFs (Figure 2B). Thus, unlike the posterior HOXC9 and HOXC10, the two main brachial central group HOX proteins bind similar target sites in differentiating neurons. To facilitate the interpretation of the subsequent comparisons, we picked the canonical HOXC6 to represent the brachial-inducing HOX binding profile.

Next, we compared the binding patterns of the central HOXC6 TF versus each of the posterior HOX TFs, HOXC9 and HOXC10. Although both induce limb-innervating spinal cord fate, HOXC6 and HOXC10 do not have identical binding patterns. While there are sites that HOXC6 and HOXC10 bind at similar levels, there are also unique HOXC6 and HOXC10 sites in differentiating cells (Figure 2C, Supp Figure 3C). Similarly, HOXC6 and HOXC9 bind to a set of sites at similar levels but there are also sites differentially bound by HOXC6 or HOXC9 (Figure 2D, Supp Figure 3D). Thus, the different patterning abilities of a central and a posterior HOX protein may be explained in part by differential genome-wide binding profiles.

To better characterize the diversity of sites bound by the various HOX TFs, we performed a joint differential binding analysis for HOXC6, HOXC9, and HOXC10 (see Methods) (Figure 2E, Supp Figure 3E). This analysis revealed that 4,892 sites were bound similarly by HOXC6, HOXC9, and HOXC10 (“**c6=c9=c10**”) (Figure 2E). HOXC6 and HOXC9 differentially bind large sets of private sites: 3,031 sites are preferentially bound by HOXC6 alone (“**c6**>c9,c10”) and 3,310 sites are preferentially bound by HOXC9 alone (“**c9**>c6,c10”). Consistent with HOXC10 binding to a subset of HOXC9 sites, fewer than 200 sites were categorized as preferentially bound by HOXC10 alone. Additionally, there are 1,106 sites preferentially bound by posterior HOX TFs (“**c9,c10**>c6”). Finally, there are 1,055 sites bound by HOXC6 and HOXC9 (“**c6,c9**>c10”). Of note, HOX binding sites are overwhelmingly distal to TSSs (Supp Figure 4A), and HOX TF binding patterns are mostly the same in newborn and young post-mitotic MNs (Supp Figure 4B-D).

Altogether, we find a set of sites that are bound by all assayed HOX TFs, regardless of their paralog group and fate inducing activity. The brachial HOXC6 and thoracic HOXC9 bind additional sets of unique sites. Finally, although the posterior HOXC10 and HOXC9 are predicted to share sequence specificity, HOXC10 cannot bind to a large fraction of HOXC9 sites.

### Sequence preference does not explain posterior HOXC9 and HOXC10 binding differences

Next, we sought to investigate whether distinct sequence preferences define the differential HOXC9 and HOXC10 binding patterns. Previous *in vitro* binding preference studies reported that anterior and central HOX paralogs prefer the TAAT core motif while the posterior paralogs prefer the TTAT motif (Mann et al., 2009; Noyes et al., 2008). Indeed, *de novo* motif discovery, using the ensemble method MEME-ChIP (Machanick & Bailey, 2011), revealed distinct motifs when comparing sites bound by central versus posterior HOX TFs during MN differentiation (Figure 3A). The representative enriched motifs detected at sites bound by HOXC6 contained the TAAT sequence. In contrast, the identified sequences at sites bound by HOXC9 and HOXC10 matched the posterior motif TTTAT and the bipartite PBX (TALE cofactor) and posterior HOX motif TGATTTAT at **c6=c9=c10** sites (Figure 3A). Thus, central and posterior HOX proteins bind to motifs in differentiating cells that are in agreement with previous *in vitro* binding preference studies (Mann et al., 2009; Noyes et al., 2008). However, both **c9**>c6,c10 and **c9,c10**>c6 binding categories have similar detected TTTAT motifs, failing to discriminate sequence preference within the posterior group. We next used motif scanning approaches to directly compare the over-representation of each type of HOX motif (the anterior TAAT and posterior TTTAT motif) at each subset of sites in the 3-way comparison. These results are consistent with the previous motif-finding results and show similar overrepresentation levels of the TTTAT motif at sites bound by HOXC9 versus HOXC9 and HOXC10 (Figure 3B).

**Figure 3:**
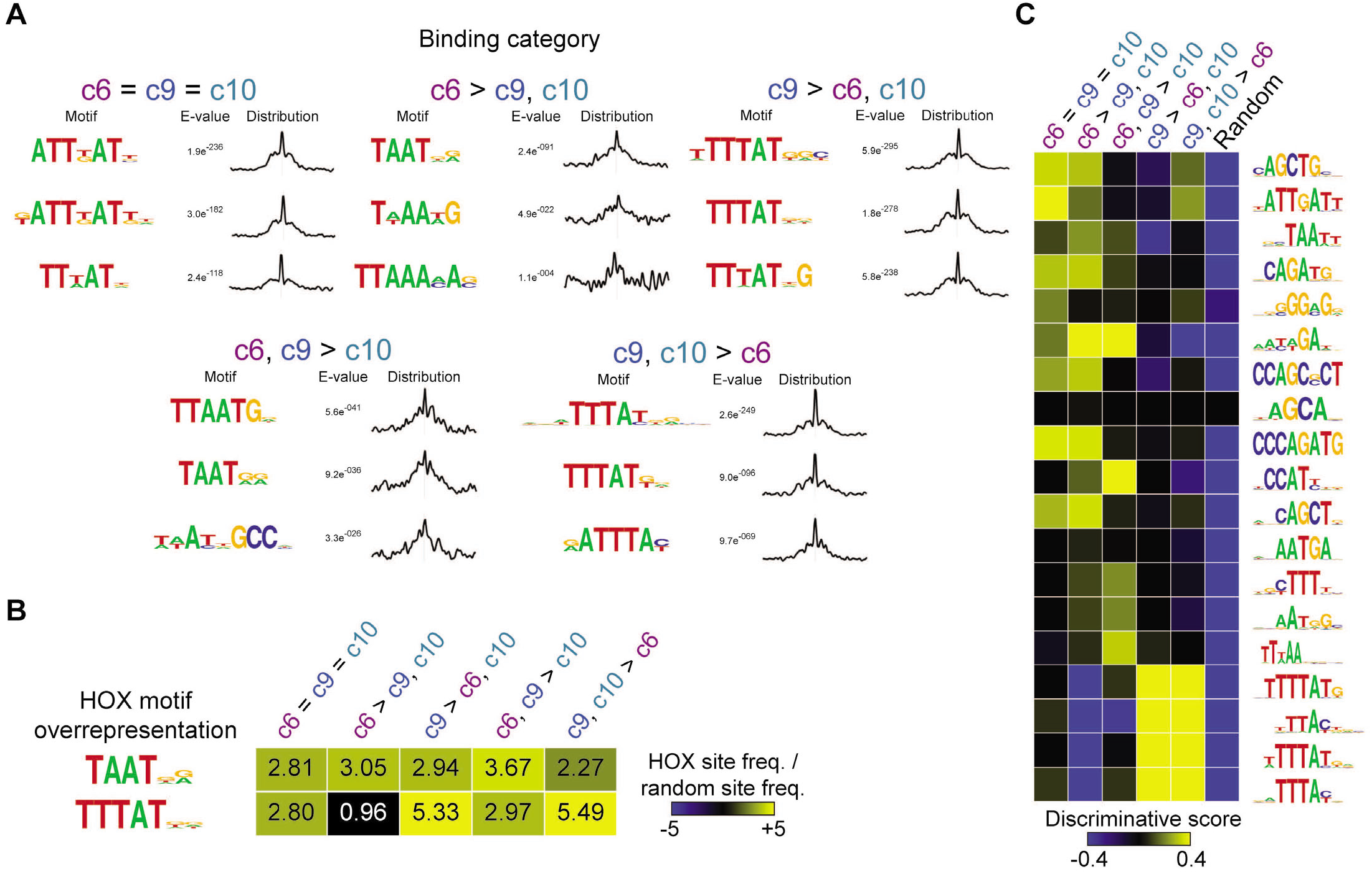
Sequence preference does not explain posterior HOXC9 and HOXC10 binding differences. (A) Selected top enriched motifs discovered via MEME-ChIP at the indicated HOX binding categories from Figure 2E. Distributions to the left of each motifs show the distribution of each motif occurrence with respect to the midpoint of each peak (500bp windows). (B) HOX TF anterior and posterior motif representation (compared with randomly selected sequences) at each category of HOX binding sites. (C) SeqUnwinder analysis characterizing motifs that are discriminative between the various classes of HOXC6, HOXC9, and HOXC10 binding sites.

Finally, we used a multi-class discriminative *k*-mer based motif-finder (SeqUnwinder) (Kakumanu, Velasco, Mazzoni, & Mahony, 2017) to find motifs that discriminate between each subset of HOX binding sites (Figure 3C). This analysis resulted in several HOX and non-HOX motifs that are specifically associated with one or more of the HOX binding site subsets (Figure 3C). For example, we find both cognate HOX TAAT motifs and additional secondary motifs that discriminate sites bound by the central HOXC6 from sites bound by the posterior HOXC9 or HOXC9 and HOXC10. Notably, however, SeqUnwinder discovers no motifs that can discriminate between sites bound by HOXC9 alone versus those bound by HOXC9 and HOXC10. Thus, we see no evidence for sequence preference differences which could explain the differential binding observed between HOXC9 and HOXC10.

### HOXC9 has a higher preference for relatively inaccessible chromatin than HOXC6 and HOXC10

In addition to a TF’s sequence preferences, cell type-specific chromatin environments can specify genomewide TF binding specificity. Since we found no strong evidence that sequence preference explains HOXC9 versus HOXC10 differences, we decided to explore whether binding to previously inaccessible sites shapes their binding patterns.

To investigate whether the chromatin accessibility landscape that exists before *Hox* induction shapes HOX TF binding patterns, we characterized genome-wide chromatin accessibility states by ATAC-seq at the progenitor stage (Figure 4A, Supp Figure 1A). The distribution of progenitor ATAC-seq read density at HOXC6, HOXC8, HOXC9, and HOXC10 sites revealed that HOXC9 binding sites had the lowest median accessibility before each factor is induced (Figure 4B). We analyzed in more detail prior accessibility differences within the different established HOX binding categories focusing on the canonical brachial (*Hoxc6)*, thoracic (*Hoxc9)* and lumbar (*Hoxc10)* genes (Figure 2E). In agreement, all sites with HOXC6 binding (**c6=c9=c10**, **c6**>c9,c10, and **c6,c9**>c10 categories) harbor similar prior accessibility landscapes (Figure 4C, D) (55%, 54%, and 45% of sites overlap accessible domains in progenitors, respectively). On the other hand, **c9**>c6,c10, and **c9,c10**>c6 binding occurs in genomic locations with much lower prior accessibility (16% and 18% of sites overlap accessible domains in progenitors, respectively) (Figure 4C, D). Interestingly, the sites with higher prior accessibility landscapes are associated with non-HOX motifs; for example, bHLH factors which are common during neuronal differentiation (Figure 3C).

**Figure 4:**
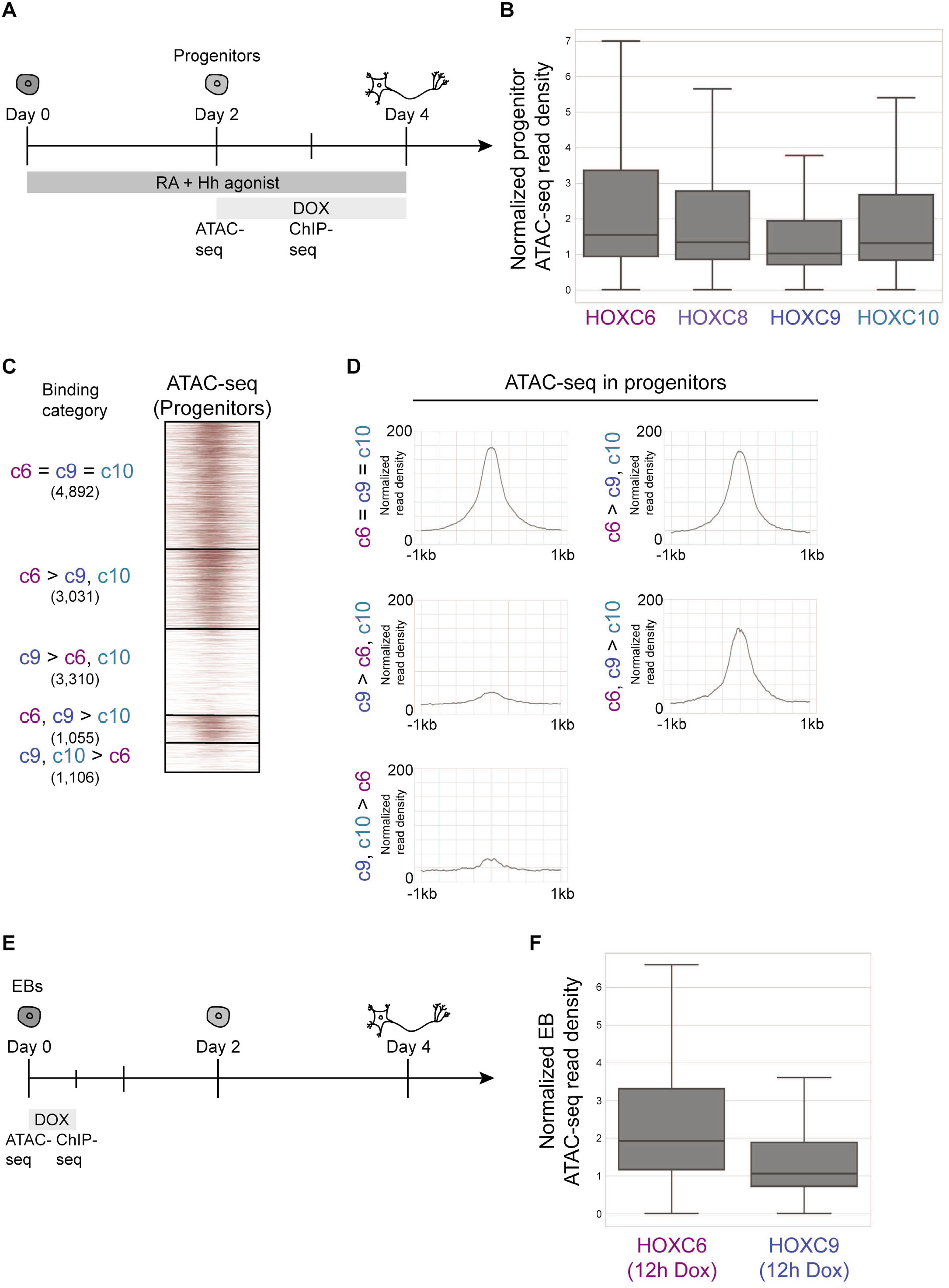
HOXC9 has a higher preference for inaccessible chromatin than HOXC6 and HOXC10. (A) Experimental scheme. (B) The distribution of Day 2 progenitor ATAC-seq read density at the top 10,000 HOXC6, HOXC8, HOXC9, and HOXC10 sites at Day 3. Data is ordered based on normalized read density (tags per million per site) and divided into quartiles. The box displays the central 50% (quartile 2 and quartile 3) of the data, while the top and bottom whiskers represent the top 25% and bottom 25% (the top and the bottom quartiles) of the data, respectively. (C) ATAC-seq heatmap displaying the accessibility in Day 2 progenitors (before HOX induction) at the indicated HOX binding categories from Figure 2E. (D) Metagene plots of accessibility (ATAC-seq reads) in progenitors displaying the prior accessibility (before HOX induction) at the indicated binding category. Normalized read density represents tags per million per 1000 sites. (E) Experimental scheme for F. (F) The distribution of EB ATAC-seq read density at the top 10,000 HOXC6 and HOXC9 sites in EBs (12h Dox induction). Data is ordered based on normalized read density (tags per million per site) and divided into quartiles. The box displays the central 50% (quartile 2 and quartile 3) of the data, while the top and bottom whiskers represent the top 25% and bottom 25% (the top and the bottom quartiles) of the data, respectively.

To test if HOX differential preferences for accessible regions were HOX TF intrinsic or due to a progenitor-specific chromatin and cofactor environment, we investigated the binding of HOXC6 and HOXC9 TFs in a different cell type (Figure 4E). Undifferentiated cells found in embryoid bodies (EBs) at an early differentiation state before the application of patterning signals have a dissimilar accessibility landscape to the neural progenitors used before. Even in this different cell type, HOXC9 maintains a higher preference for inaccessible chromatin than HOXC6 (Figure 4F). Thus, the intrinsic ability of HOX TFs to bind to inaccessible chromatin seems to be independent of the particular cellular environment in which they are expressed.

Altogether, comparing HOX TF binding with prior accessibility revealed that limb-innervating HOXC10 and HOXC6 rely more on chromatin accessibility established at progenitor stages to find their target sites. Moreover, these results divide the posterior paralog group by their ability to bind previously inaccessible chromatin, with HOXC9 displaying a greater ability to bind inaccessible chromatin when compared to HOXC10.

### HOX TF binding increases chromatin accessibility

The difference in the abilities of HOX TFs to bind to inaccessible chromatin prompted us to ask whether HOX TFs change the accessibility landscape after binding. Would HOXC9’s ability to bind inaccessible sites be coupled with increasing accessibility after binding? To characterize the accessibility changes after HOX binding, we compared the accessibility status of progenitors and post-mitotic neurons at HOXC6, HOXC9, and HOXC10 binding events (Figure 5A, Supp Figure 1A).

**Figure 5:**
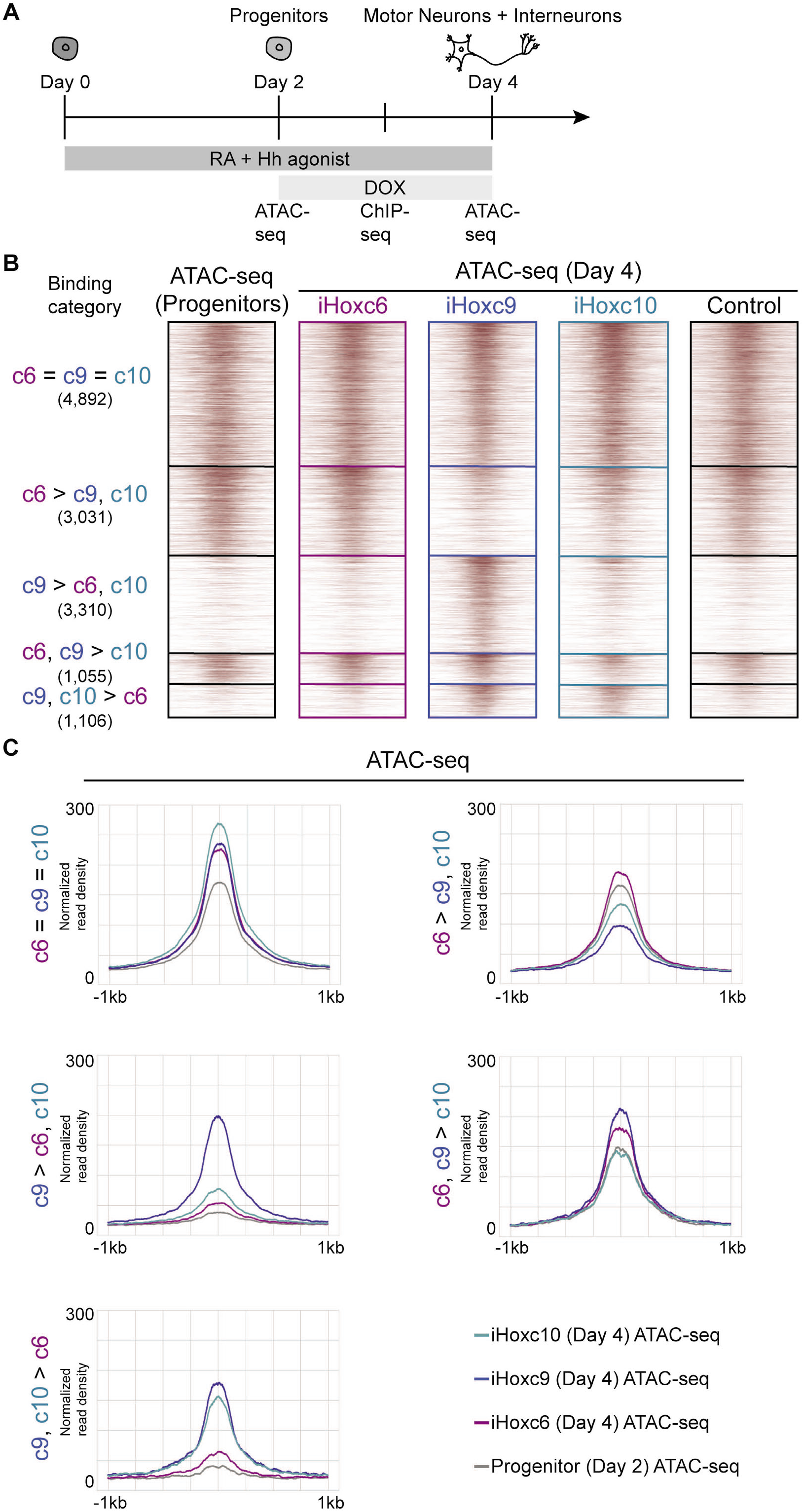
HOX TF binding increases chromatin accessibility. (A) Experimental scheme. (B) ATAC-seq heatmaps displaying the accessibility in Day 2 progenitors versus iHoxc6 versus iHoxc9 versus iHoxc10 versus No Dox control neurons, at the indicated binding categories from Figure 2E. (C) Metagene plots of accessibility (ATAC-seq reads) showing the accessibility gain in iHoxc6 versus iHoxc9 versus iHoxc10 neurons at the indicated binding category. Normalized read density represents tags per million per 1000 sites.

We find that sites bound by HOXC9 and HOXC10 (**c9**, **c10**>c6) gained accessibility in both iHoxc9 and iHoxc10 neurons, while not significantly changing in iHoxc6 cells (Figure 5B, C). Strikingly, exclusive HOXC9 sites increased accessibility the most and only in response to *Hoxc9* expression (Figure 5B, C). Consistent with HOX TFs maintaining the preexisting chromatin status, **c6**=**c9**=**c10** sites gained some accessibility after HOXC6, HOXC9, and HOXC10 binding (Figure 5B, C). Accordingly, **c6**>c9,c10 sites gained some accessibility in iHoxc6 neurons, while they lost accessibility in iHoxc9 and iHoxc10 expressing cells.

The dynamic accessibility changes after HOX binding revealed that not all HOX TFs have the same ability to modify chromatin accessibility. Among those we analyzed, HOXC9 stands out in its ability to bind to a large set of sites in inaccessible chromatin and increase the accessibility status after binding.

### Posterior group HOX TFs display a range of abilities to bind inaccessible chromatin

We wondered next if the ability of HOXC9 to bind to a large set of previously inaccessible sites is unique among posterior HOX TFs. Thus, we compared the binding of HOX9 paralog group TFs. A comparison of induced HOXA9 and HOXD9 ChIP-seq using MultiGPS revealed that they share a majority of their binding sites (Figure 6A). However, comparing HOXC9 and HOXA9 binding patterns showed that HOXC9 uniquely binds an additional large category of sites (Figure 6B). Suggesting a shared sequence preference, the detected motifs at sites bound by HOXA9 and HOXC9 resemble the posterior HOX TTTAT motif (Figure 6C). HOXC9 binds to sites with the lowest median prior accessibility among the HOX9 paralogs (Figure 6D). Accordingly, the sites uniquely bound by HOXC9 showed lower prior accessibility than other sets of sites (Figure 6E, F). Hence, our results suggest that there is a divergence in the ability to bind inaccessible sites, even within the HOX9 posterior paralog group.

**Figure 6:**
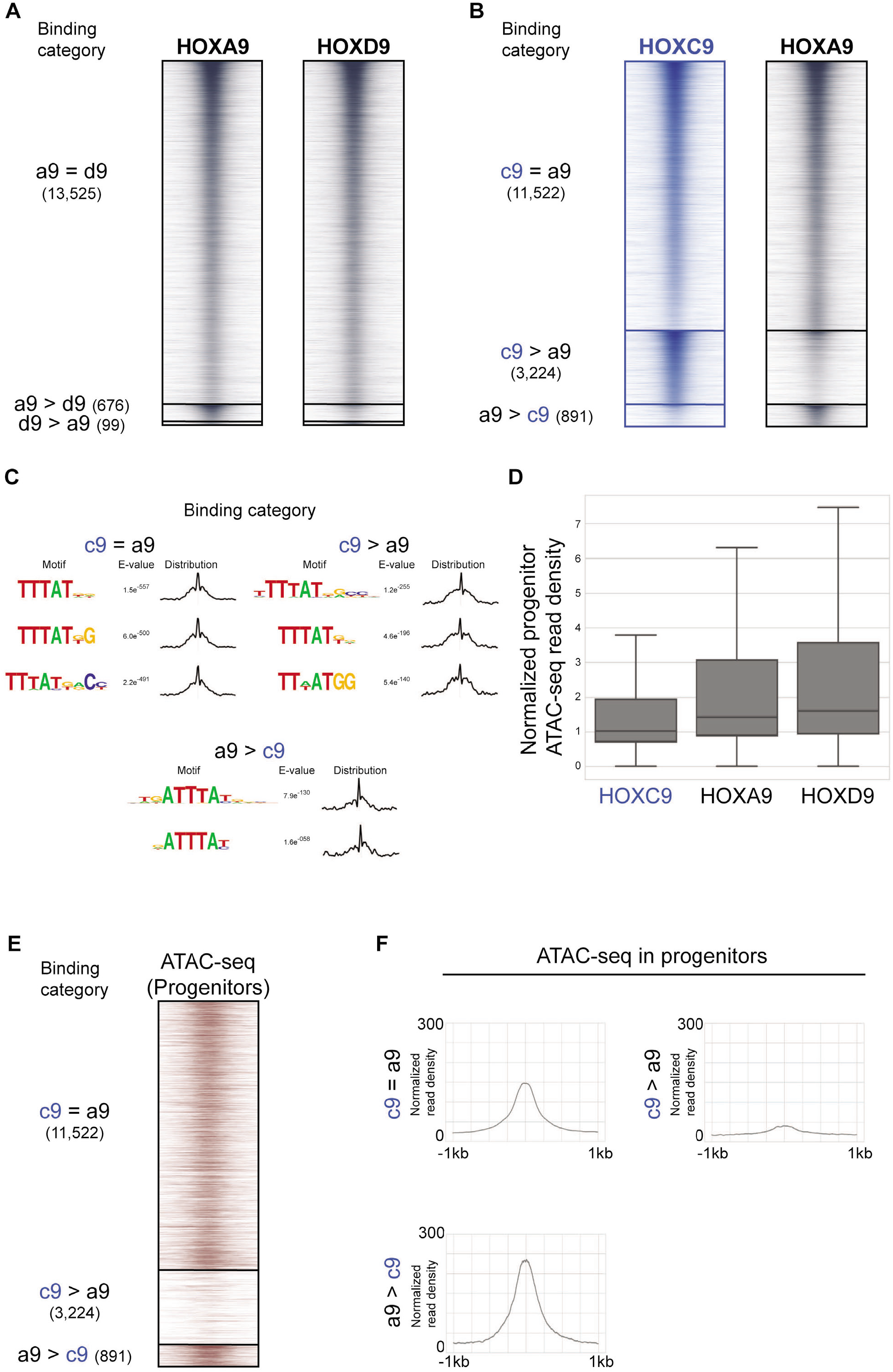
HOXC9 binds to a larger fraction of sites in inaccessible chromatin than HOXA9 and HOXD9. (A-B) ChIP-seq heatmap showing binding comparisons of the indicated HOX TFs in differentiating neurons, at Day 3. Sites bound by both indicated HOX TFs noted as “=” sites. Preferentially bound sites by HOXA9, HOXD9 or HOXC9 noted as “a9 >”, “d9 >” or “c9 >”. (C) Selected top enriched motifs discovered via MEMEChIP at the indicated HOX binding categories. Distributions to the left of each motifs show the distribution of each motif occurrence with respect to the midpoint of each peak (500bp windows). (D) The distribution of Day 2 progenitor ATAC-seq read density at the top 10,000 HOXC9, HOXA9, and HOXD9 sites at Day 3. Data is ordered based on normalized read density (tags per million per site) and divided into quartiles. The box displays the central 50% (quartile 2 and quartile 3) of the data, while the top and bottom whiskers represent the top 25% and bottom 25% (the top and the bottom quartiles) of the data, respectively. (E) ATAC-seq heatmap displaying the accessibility in Day 2 progenitors (before HOX induction) at the indicated HOX binding categories. (F) Metagene plots of accessibility (ATAC-seq reads) in progenitors displaying the prior accessibility (before HOX induction) at the indicated binding categories. Normalized read density represents tags per million per 1000 sites.

The posterior HOX binding profiles we have analyzed revealed that they do not entirely overlap in their genomic binding, despite sharing motif preferences. Thus, we sought to expand these analyses to another relevant posterior *Hox* gene. *Hox13* paralogs are also posterior group genes but have the unique ability to terminate axial elongation. To assess whether HOXC13 would bind like the other posterior HOX TFs, we compared HOXC9, HOXC10, and HOXC13 genomic binding overlaps (Figure 7A). Surprisingly, only a small fraction of all sites (1,024) are shared by HOXC9, HOXC10, and HOXC13 (“**c9=c10=c13**”). As in all previous comparisons, HOXC9 retained a subset of private sites 2,543 (“**c9**>c10,c13”). However, in this comparison, a large category (5,652) of sites was bound by HOXC13 alone (“**c13**>c9,c10”). A direct comparison of HOXC9 and HOXC13 binding profiles supported the finding that they primarily bind distinct sets of sites (Supp Figure 5A). Motif analysis at these sites revealed that HOXC13 binds distinct motifs containing the TTTAC sequence (Supp Figure 5B, C), in agreement with previous *in vitro* binding characterizations (Berger et al., 2008). Thus, HOXC13 has a distinct motif preference, thereby increasing the posterior HOX TF binding diversity.

**Figure 7:**
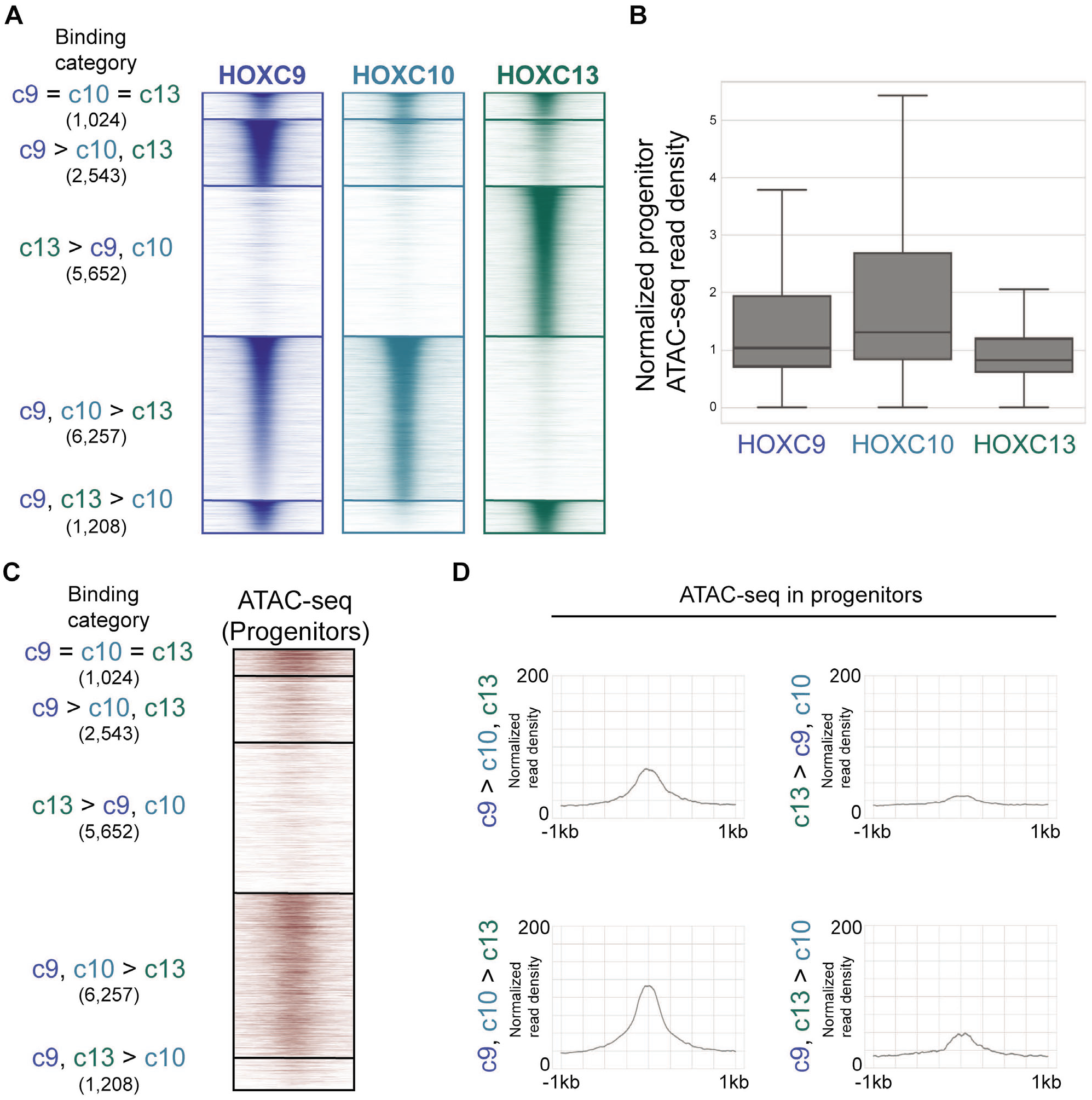
Posterior group HOX TFs display a range of abilities to bind inaccessible chromatin. (A) ChIP-seq heatmap showing binding comparisons of HOXC9, HOXC10, and HOXC13 in differentiating neurons, at Day 3. Sites bound by all three HOX TFs noted as “c9 = c10 = c13” sites. Preferentially bound sites by HOXC9, HOXC13, HOXC9 and HOXC10 or HOXC9 and HOXC13 noted as “c9 > c10, c13”, “c13 > c9, c10”, “c9, c10 > c13”, and “c9, c13 > c10”, respectively. (B) The distribution of Day 2 progenitor ATAC-seq read density at the top 10,000 HOXC9, HOXC10, and HOXC13 sites at Day 3. Data is ordered based on normalized read density (tags per million per site) and divided into quartiles. The box displays the central 50% (quartile 2 and quartile 3) of the data, while the top and bottom whiskers represent the top 25% and bottom 25% (the top and the bottom quartiles) of the data, respectively. (C) ATAC-seq heatmap displaying the accessibility in Day 2 progenitors (before HOX induction) at the indicated HOX binding categories. (D) Metagene plots of accessibility (ATAC-seq reads) in progenitors displaying the prior accessibility (before HOX induction) at the indicated binding categories. Normalized read density represents tags per million per 1000 sites.

We then asked if the posterior HOX group TF binding diversity also correlates with their differential ability to bind to previously inaccessible chromatin. The distribution of progenitor ATAC-seq read density at HOXC9, HOXC10, and HOXC13 sites revealed that HOXC13 binding sites had the lowest median accessibility in MN progenitors, even lower than HOXC9 (Figure 7B). Dissecting the accessibility at the different HOX binding categories underscored the ability of HOXC13 to bind to sites with the lowest prior accessibility (Figure 7C, D). In agreement, a recent *in vivo* study revealed that the patterning activity of HOX13 paralogs during limb development relies on their ability to increase accessibility at specific sites (Desanlis et al., 2020).

To visualize the overall variation in HOX TF genome-wide binding profiles, we performed Principal Components Analysis (PCA) on HOX TF ChIP-seq read counts associated with the top 10,000 binding sites for at least one HOX TF (38,874 unique genomic locations; Figure 8A, Supp Figure 5D, E). PC1, which explains 43% of the variance between the TFs, separates HOXC13 from the central HOX TFs, HOX9 paralogs, and HOXC10. On the other hand, PC2 and PC3, which cumulatively explain 32% of the variance, separate the TFs into three clusters. Although distinguishable, HOXC6 and HOXC8 cluster close to each other. HOXC9 clusters by itself, and finally, HOXC10 and HOX9 paralogs HOXA9 and HOXD9 cluster together.

**Figure 8:**
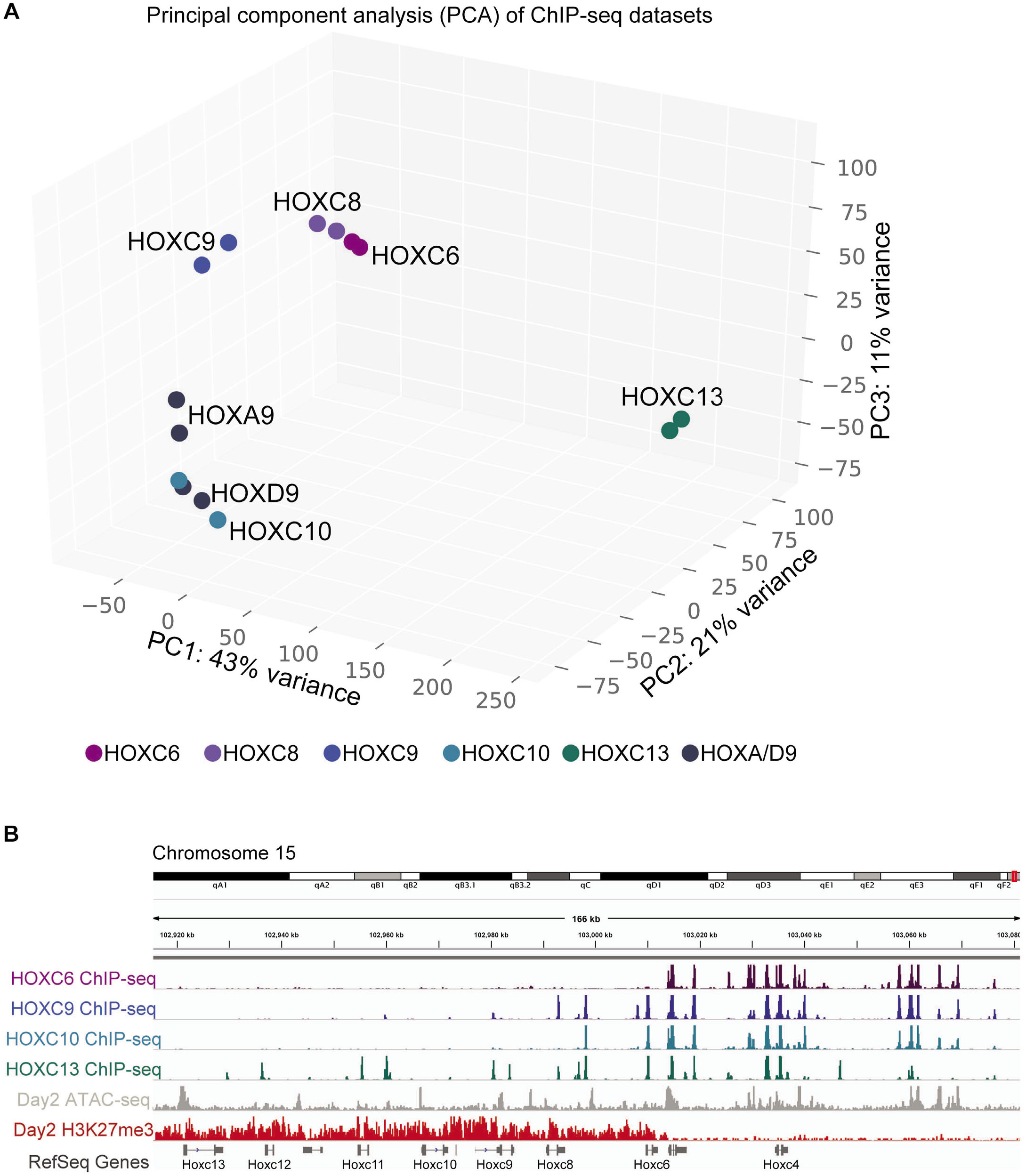
Posterior HOX TFs have distinct genomic binding patterns. (A) Principal Component Analysis (PCA) of the ChIP-seq datasets reveals similarities in the binding patterns of HOX TFs. (B) Browser screenshots of the indicated HOX ChIP-seqs, Day 2 ATAC-seq and H3K27me3 ChIP-seq at the *HoxC* gene cluster.

The binding pattern of HOXC TFs to the *HoxC* gene cluster demonstrates their differential ability to engage inaccessible chromatin (Figure 8B). While HOXC6 only binds *Hoxc4-5* genes, which are transcriptionally active, HOXC10, HOXC9, and HOXC13 bind progressively deeper into the cluster at repressed *HoxC* genes which are covered in the catalytic product of PRC2 (Mazzoni et al., 2013; Narendra et al., 2015). Thus, differences in genome-wide binding profiles of the HOX TFs reflect sequence preferences, as well as the differential abilities of the HOX TFs to bind previously inaccessible chromatin. Posterior HOX TFs can bind different genomic sites by either having similar binding preferences and differing abilities to bind inaccessible chromatin (HOXC9 vs. HOXC10), or differing in both aspects (HOXC9 vs. HOXC13).

Finally, to gain an overview of each HOX TF’s relative dependence on the preexisting chromatin environment, we applied Bichrom to analyze the data (Srivastava, Aydin, Mazzoni, & Mahony, 2019). Bichrom is a neural network-based method that integrates DNA sequence and prior chromatin information to explain the observed genomic binding patterns of an induced TF. Specifically, we train Bichrom to predict the binding patterns of each HOX TF using DNA sequence features and a selection of chromatin tracks from Day 2 progenitors (ATAC-seq and ChIP-seq for H3K4me3, H3K27ac, H3K27me3, H3K9me3, and Pol II). We then compared Bichrom’s predictive performance with that of a baseline convolutional and long short-term memory neural network (CNN-LSTM) that uses only sequence information (seqnet) (Supp Figure 6A). We trained both networks using 9 distinct training sets (each associated with an independent held-out test set). Following training, we compared the performance of the sequence-only network to that of Bichrom on held-out test sets using the area under the precision-recall curve (auPRC). If all HOX TFs had similar reliance on chromatin states for binding, we would predict similar recall improvements across TFs when incorporating chromatin data in addition to the sequence. However, our data shows a variation that correlates with their preference for inaccessible chromatin (Figure 4B, 6D, 7B). We find that HOXC9 and HOXC13 networks display minor improvements in the predictive performance when trained with or without neuronal progenitor chromatin data. Consistent with our previous results, none of the various incorporated prior chromatin tracks predict future binding for those posterior TFs. HOXC8 and HOXC10 predictions benefit from included prior chromatin data, but HOXC6 and the other HOX9 paralogs (HOXA9 and HOXD9) display substantial gains in predictive performance when training includes preceding chromatin tracks. These results support the hypothesis that even HOX TFs from the same group rely on previous chromatin states to different degrees for their genomic binding.

### The genomic binding of HOXC6, HOXC9, and HOXC10 correlates with differential gene expression

Finally, we asked whether differentially expressed genes in the iHox neurons correlate with specific HOX binding categories. For this comparison, we focused on the three main spinal cord domains and their “canonical” inducing TFs: HOXC6 for brachial, HOXC9 for thoracic, and HOXC10 for lumbar. Specifically, we used the logistic regression-based ChIP-Enrich method to identify significant associations (adjusted p-value < 0.01) between RNA-seq derived gene sets and HOX TF binding categories. Genes that are equally up-regulated in iHoxc6, iHoxc9, and iHoxc10 neurons associated strongly with **c6=c9=c10** sites, with some association with **c6**>c9,c10 and **c9,c10**>c6 sites as well (Figure 9A). Genes differentially expressed in iHoxc6, compared to iHoxc9 or iHoxc10 neurons, show a correlation with **c6**>c9,c10 sites (Figure 9A). Similarly, genes differentially expressed in iHoxc9 neurons correlate with **c9**>c6,c10 sites (Figure 9A). Thus, HOX TF binding correlates with transcriptional activity. Moreover, the enriched GO terms at HOX binding sites are relevant to the phenotypes induced by the HOX TFs *in vivo* (Figure 9B-F), even with all the limitations of GO term analysis for tissue segments (spinal cord regions). For example, neuron differentiation and central nervous system development appear as the top GO terms at several binding categories. Also, several GO terms on axon development and guidance appear at sites preferentially bound by HOXC9 (Figure 9D), which are critical motor neuron features well documented to be downstream of HOX patterning.

**Figure 9:**
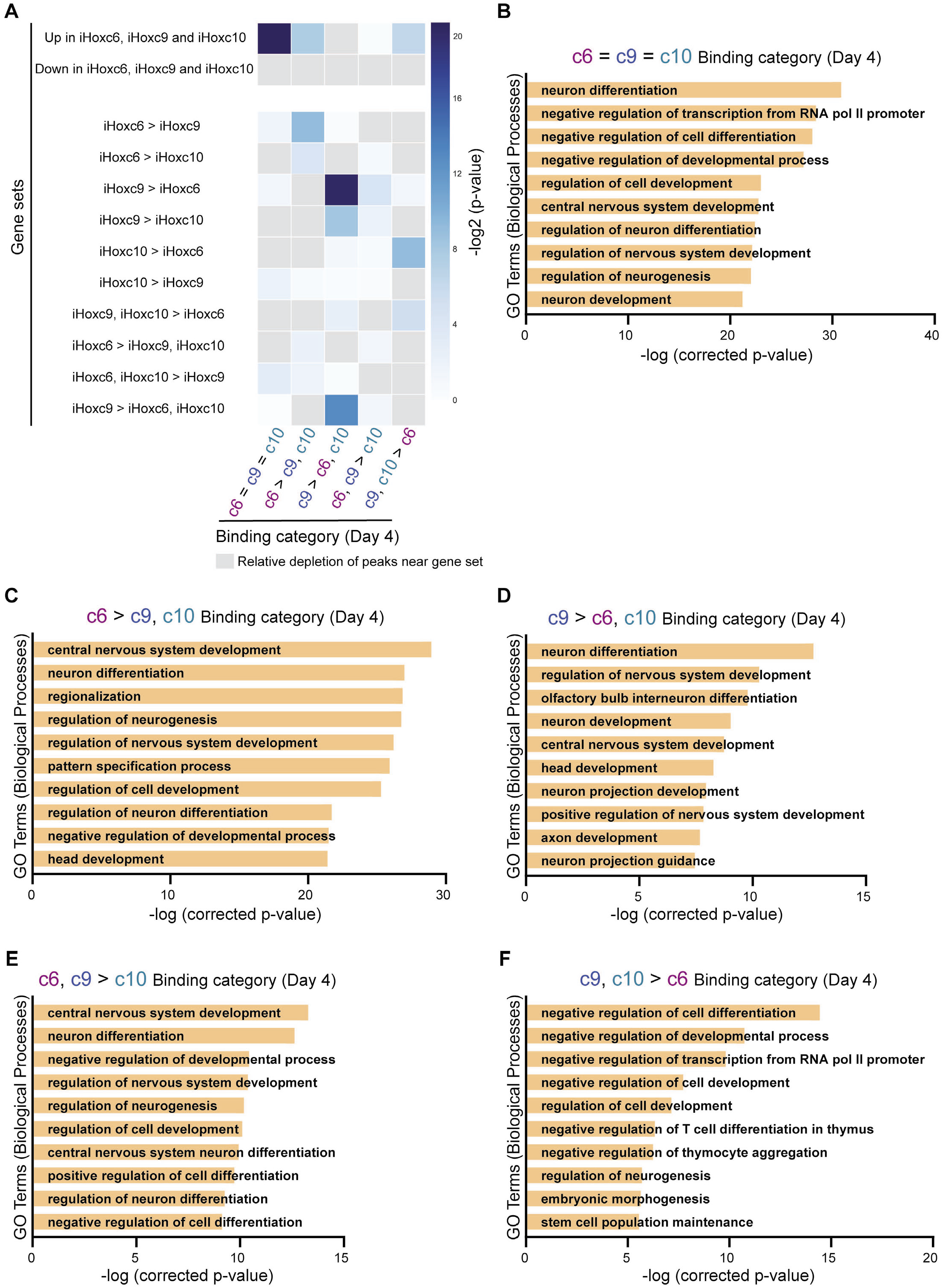
Differentially expressed gene sets correlate with differential HOX binding events. (A) Heatmap representing associations between HOX binding categories (Day 4; from Supp Figure 3E) and the indicated gene sets (Day 4). Upregulated/Downregulated in iHoxc6, iHoxc9, and iHoxc10 (Day 4) is relative to Day 2 progenitors (prior to HOX induction). (B-F) Top GO-terms enriched at the indicated HOX binding categories (Day 4; from Supp Figure 3E).

## DISCUSSION

HOX TFs have crucial roles in body patterning during animal development. However, surprisingly little is known about how vertebrate HOX TFs bind to the genome in a cellular-relevant environment. This is exacerbated for posterior group *Hox* genes that pattern distinct vertebrate structures, albeit sharing a common *Drosophila* ortholog gene. To gain insights into HOX activity, we performed a multilevel comparison of global binding patterns, chromatin accessibility preferences, and transcriptional target genes of seven HOX proteins expressed under the same developmentally relevant conditions. While the data show intrinsic sequence preferences that differ between HOX TFs, we find that a major determinant of genomic binding diversity amongst posterior HOX TFs are their differential abilities to bind inaccessible chromatin. Therefore, posterior HOX TF patterning may diverge by mostly tuning chromatin accessibility binding rather than sequence preference.

The central group HOXC6 and HOXC8 TFs induce limb-innervating fate at brachial spinal cord levels. They appear to do so by binding to relatively similar sites in the genome compared to other analyzed HOX binding profiles. HOXC10 induces a similar limb-innervating fate, albeit at lumbar levels, but it has a binding profile more similar to the thoracic HOXC9 (repressor of limb-level fates) than to HOXC6 or HOXC8. Overall, these results suggest that similar cell fates are not always the product of identical HOX TF binding patterns. HOXC10 sequence preference is in line with the posterior group. However, HOXC10’s ability to bind to inaccessible chromatin is more similar to central limb-innervating HOXC6 and HOXC8. Thus, HOXC10 diverges from HOXC9 by having a lower preference for inaccessible chromatin.

During embryonic development, limb-innervating LMC neurons are further patterned by HOX proteins into specific pools (Dasen, Tice, Brenner-Morton, & Jessell, 2005). During this second stage, HOXC6 and HOXC8 TFs differ in their fate-inducing identity. The *in vitro* differentiation strategy and the forced expression of a single *Hox* gene does not recapitulate this late differentiation event. Thus, we speculate that pool patterning by *Hox* genes does require a more complex developmental context expressing more *Hox* genes and signaling molecules important for pool identity acquisition (Arber, Ladle, Lin, Frank, & Jessell, 2000; Dasen et al., 2005; Haase et al., 2002). Although technically unreachable for now, future HOX binding in individual brachial motor neuron pools may elucidate how HOXC6 and HOXC8 binding induces different motor neuron pool subtypes. Additionally, *Hoxc10* overexpression induces some *Hoxc6* expression (Figure 1D, Supp Figure 2A-C). It will be interesting to test if an *in vivo Hoxc10* overexpression induces additional limb-innervating *Hox* genes.

In agreement with *in vitro* binding preference studies, central and posterior HOX proteins bind to different motifs in neurons (Berger et al., 2008; Mann et al., 2009; Noyes et al., 2008). While central HOX TFs bind to sites with the central TAAT core motif, posterior HOX TFs bind to TTAT core motifs. MEME-ChIP also discovered bipartite cofactor and HOX motifs with high enrichment in some sets of sites (Figure 3A). An interaction with TALE cofactors, MEIS and PBX, can change the affinity and selectivity of HOX DNA-binding (Slattery et al., 2011) (reviewed in (Mann & Affolter, 1998; Mann & Chan, 1996; Merabet & Mann, 2016)). Besides, partnering with TALE cofactors modifies HOX binding specificity through the recognition of DNA shape (Abe et al., 2015; Joshi et al., 2007; Zeiske et al., 2018). Although *Hox* expression changes the fate of *in vitro* differentiating neurons, differential binding of the canonical MEIS HOX cofactor does not seem to explain differential HOX TF binding patterns (Supp Figure 7A). MEIS binding appears to follow or reflect HOX binding, as opposed to being exclusively associated with particular subsets of differentially bound sites.

Furthermore, PBX1-4 and MEIS1-3 expression levels are largely similar across the different HOX TF inductions and are thus unlikely to explain binding differences (Figure 1D, Supp Figure 2A, B). We attempted similar experiments with PBX factors in two of the inducible *Hox* lines, but the PBX antibody produces a weak ChIP-seq signal (Supp Figure 7B). We also failed to detect a CTCF motif at HOX binding sites, which is reported to co-bind with some HOXA/D proteins (Jerkovic et al., 2017). Our data thus failed to identify a cofactor that explains differential HOX binding during motor neuron differentiation. A systematic evaluation and perturbation of all possible HOX cofactors during cell differentiation will shed some light on this issue.

Our data suggest HOX binding, and thus patterning, models should integrate sequence and chromatin state to explain each HOX activity. Most HOX TFs would bind to cell-specific accessible sites with canonical motifs. Hence, HOXC6, HOXC9, and HOXC10 shared sites tend to be in regions with high prior accessibility during motor neuron differentiation, and these sites appear to contain both types of HOX binding motifs. However, some HOX TFs can associate with sites in less accessible genomic regions. Thus, they become more independent of earlier chromatin patterning events. In sum, the data presented here suggest HOX TF patterning abilities can only be explained by integrating not just each HOX TF’s sequence preference and cofactor interactions, but also the preexisting cell-specific chromatin landscape and the HOX TF’s ability to interact with inaccessible chromatin.

## DATA AVAILABILITY

Sequencing data have been submitted to the GEO under accession code GSE142379.

## ACKNOWLEDGEMENTS

This work was supported by the NICHD (R01HD079682) and NINDS (R01NS100897) grants to E.O.M, the NYSTEM pre-doctoral training grant (C322560GG) to M.B, the Penn State Academic Computing Fellowship to D.S, the NINDS (R01NS062822) grant to J.S.D and NIGMS (R01GM121613) to S.M. The authors would like to thank Ayana Sawai-Frantz for sharing the pCAGGS mHoxA9 (JD-114) and pCAGGS mHoxD9 (JD-237) plasmids; the Mazzoni and Mahony lab members for feedback and constructive comments; the NYU Genomics Core facility for technical support and data processing.

## AUTHOR CONTRIBUTIONS

M.B performed *in vitro* differentiation, ChIP-seq, RNA-seq, and ATAC-seq experiments and established inducible cell lines. D.S performed the analysis of sequencing data with help from M.B and S.M. J.S.D and H.W provided plasmids, cell lines, and guidance. M.B, D.S, S.M, and E.O.M conceived the experiments and set up the computational analysis framework. M.B and E.O.M co-wrote the original draft of the manuscript. All authors read and approved the final manuscript.

## DECLARATION OF INTEREST

Authors declare no competing interests.

## METHODS

### Cell line generation

The inducible Hoxc6, Hoxc8, and Hoxc9 cell lines were generated as previously described (Mazzoni et al., 2011). The inducible cassette exchange (ICE) system was used to generate cell lines (Iacovino et al., 2011). The resulting inducible cell lines harbor a single copy of the transgene inserted at the expression-competent HPRT locus. Inducible Hoxa9, Hoxd9, Hoxc10, and Hoxc13 cell lines were generated for this study. Hoxa9, Hoxd9, and Hoxc13 cDNA was amplified using Phusion polymerase (Thermo Scientific) from pCAGGS mHoxA9 (JD-114), pCAGGS mHoxD9 (JD-237) and hHoxc13 cDNA (Dharmacon, Accession: BC090850), respectively. Flag and HA tags were introduced during the amplification step at the amino- or carboxyl-terminus, respectively. p2lox plasmids were generated by In-Fusion cloning (Takara) the respective cDNAs into the p2lox plasmid backbone. The recipient mESCs were treated with 1 μg/mL Doxycycline (Sigma, D9891) for 16h to induce Cre recombinase expression prior to electroporation of the respective plasmids. After selection with G418 (250ng/mL, Cellgro), cell lines were characterized by performing antibody staining for Flag (mouse anti-FLAG; Sigma, F1804) and HA (rabbit anti-HA; Abcam, ab9110) and expanded.

### Cell culture

mESC lines were cultured in 2-inhibitors based medium (Advanced DMEM/F12:Neurobasal (1:1) medium (Gibco), supplemented with 2.5% ESC-grade fetal bovine serum (vol/vol, Corning), N2 (Gibco), B27 (Gibco), 2mM L-glutamine (Gibco), 0.1 mM ß-mercaptoethanol (Gibco), 1000 U/mL leukemia inhibitory factor (Millipore), 3μM CHIR (BioVision) and 1 μM PD0325901 (Sigma)) on 0.1% gelatin (Millipore) coated plates at 37 °C, 8% CO2.

*In vitro* differentiation of mESCs to motor neurons was described previously (Tan et al., 2016; Wichterle et al., 2002; Wichterle & Peljto, 2008). Briefly, embryoid bodies (EBs) were obtained by plating trypsinized (Gibco) mESCs in AK medium (Advanced DMEM/F12:Neurobasal (1:1) medium (Gibco), 7% KnockOut SR (vol/vol) (Gibco), 2mM L-glutamine, 0.1 mM ß-mercaptoethanol and penicillin–streptomycin (Gibco),) at 37°C, 5%CO2 (day −2). On day 0, EBs were split 1:2 and AK medium was replenished and supplemented with 1 μM all-trans retinoic acid and 0.5 μM smoothened agonist (SAG) (Millipore, 566660). TF induction was performed by adding 3 μg/mL of Doxycycline (Sigma, D9891) on day 2. For RNA-seq and ATAC-seq experiments, 3.5×10^5^ mESCs were plated in 100mm suspension dishes (Corning). For ChIP-seq experiments, 3-3.5×10^6^ mESCs were plated in 245mm x 245mm square dishes (Corning).

### ChIP-seq

Cells were collected 24h and 48h after Doxycycline (Dox) treatment (day 3 and 4 of RA/SAG differentiation). Crosslinking was performed at room temperature in 1mM DSG (ProteoChem) for 15min, followed by the addition of 1% FA (vol/vol) for an additional 15min. After quenching with Glycine, cells were divided into ~25-30×10^6^ aliquots, pelleted by centrifugation and frozen at −80°C. After thawing cells on ice, lysis was performed in 5mL of 50 mM HEPES-KOH pH7.5, 140 mM NaCl, 1 mM EDTA pH 8.0, 10% glycerol (vol/vol), 0.5% Igepal (vol/vol), 0.25% Triton X-100 (vol/vol) with 1× protease inhibitors (Roche, 11697498001) for 10 min at4°C. Cells were centrifuged for 5min, resuspended in 5mL of 10 mM Tris-HCl pH 8.0, 200 mM NaCl, 1 mM EDTA pH 8.0, 0.5 mM EGTA pH 8.0 with 1× protease inhibitors, and incubated for 10 min at 4 °C on a rotating platform. Cells were centrifuged for 5min and resuspended in 2mL of Sonication Buffer (50 mM Hepes pH 7.5, 140 mM NaCl, 1 mM EDTA pH 8.0, 1 mM EGTA pH 8.0, 1% Triton X-100 (vol/vol), 0.1% sodium deoxycholate (wt/vol), 0.1% SDS (vol/vol) with 1× protease inhibitors). Sonication was performed by splitting each sample to 2 Bioruptor tubes with sonication beads and using the Bioruptor Pico (Diagenode) for 18 cycles of 30 sec on and 30 sec off into an average size of approximately 200 bp. Immunoprecipitation was performed for 16h at 4°C on a rotating platform by incubating with Dynabeads protein-G (Thermo Fisher Scientific) conjugated with 5 μg of one of the following antibodies: mouse monoclonal antibody to Flag (Sigma, F1804) or rabbit polyclonal antibody to HA (Abcam, ab9110). Following the immunoprecipitation, washes were performed with the following buffers (cold): sonication buffer, sonication buffer with 500 mM NaCl, LiCI wash buffer (20 mM Tris-HCI pH 8.0, 1 mM EDTA pH 8.0, 250 mM LiCI, 0.5% Igepal (vol/vol), 0.5% sodium deoxycholate (wt/vol)), and TE buffer (10 mM Tris-HCl pH 8.0, 1 mM EDTA pH 8). Elution was performed by incubating in elution buffer (50 mM Tris-HCI pH8.0, 10mM EDTA pH 8.0, 1% SDS (vol/vol)) for 45 min at 65 °C. Reversal of crosslinks was performed by incubating for 16h at 65 °C. RNA digestion was performed by the addition of 200 μL of TE and RNAse A (Sigma) at a final concentration of 0.2 mg/mL and incubation for 2 h at 37°C. Proteinase K (Invitrogen) was added at a final concentration of 0.2 mg/mL, supplemented with CaCl2, to digest protein at 55°C for 30 min. DNA was purified with phenol:chloroform:isoamyl alcohol (25:24:1; vol/vol) (Invitrogen) and by performing ethanol precipitation. DNA pellets were resuspended in water. lllumina DNA sequencing libraries were prepared with one third of the ChIP sample or a 1:100 dilution of the input sample in water. Library preparation was performed by end repair, A-tailing and ligating Illumina-compatible Bioo Scientific multiplexed adapters. Unligated adapters were removed using Agencourt AmpureXP beads (Beckman Coulter). Amplification was performed by PCR with Phusion polymerase (New England Biolabs) and TruSeq primers (Sigma). Libraries were gel purified (Qiagen) between 250 and 550bp in size. Final quantification of the library was performed using the KAPA library amplification kit on the Roche Lightcycler 480 before pooling. The libraries were sequenced on Illumina NextSeq 500 using V2 and V2.5 chemistry (75 cycles, single-end 75bp) at the Genomics Core Facility at NYU.

### RNA-seq

Cells were collected prior to TF induction (day 2 of RA/SAG differentiation) and 48h after Doxycycline (Dox) treatment (day 4 of RA/SAG differentiation). RNA was extracted by using TRIzol LS Reagent (Life Technologies) and purified using the RNAeasy mini kit (Qiagen). Agilent High Sensitivity RNA Screentape (Agilent, 5067-5579) was used to check RNA integrity. 500ng of RNA was used to prepare RNA-seq libraries and spiked-in with ERCC Exfold Spike-in mixes (Thermo Fisher, 4456739). RNA-seq libraries were prepared using TruSeq Stranded mRNA Library Preparation kit (Illumina, 20020594). Library size was verified using High Sensitivity DNA ScreenTape (Agilent, 5067-5584). The KAPA library amplification kit was used on Roche Lightcycler 480 for library quantification before pooling. The libraries were sequenced on Illumina NextSeq 500 using V2.5 chemistry (75 cycles, single-end 75bp) at the Genomics Core Facility at NYU.

### ATAC-seq

Cells were collected prior to TF induction (day 2 of RA/SAG differentiation) and 48h after Doxycycline (Dox) treatment (day 4 of RA/SAG differentiation). 50,000 cells were aliquoted and washed twice in 1 × PBS (cold). Cell pellets were resuspended in 10 mM Tris pH 7.4, 10 mM NaCl, 3 mM MgCl_2_, and freshly added 0.1% NP-40 (vol/vol), and centrifuged at 4 °C. Pellets were resuspended in 25 μL of 2 × TD buffer, 2.5 μL TDE1 (Nextera DNA sample preparation kit, FC-121–1030) and 22.5 μL of water and incubated at 37 °C for 30 min. The sample was purified using the MinElute PCR purification kit (Qiagen, 28004). A quantitative PCR reaction with 1 × SYBR Green (Invitrogen), custom-designed primers and 2 × NEB MasterMix (New England Labs, M0541) was performed to determine the optimal number of PCR cycles (one-third of the maximum measured fluorescence) (Buenrostro, Giresi, Zaba, Chang, & Greenleaf, 2013). PCR enrichment of the library was performed with custom-designed primers and 2 × NEB MasterMix (New England Labs, M0541). The libraries were purified using the MinElute PCR purification kit. High Sensitivity DNA ScreenTape (Agilent, 5067-5584) was used to verify the fragment length distribution of the library. Library quantification was performed using the KAPA library amplification kit on the Roche Lightcycler 480. The libraries were sequenced on Illumina NextSeq 500 using V2 and V2.5 chemistry (150 cycles, paired-end 75bp) at the Genomics Core Facility at NYU.

### ChIP-seq Data Processing

ChIP-seq reads were aligned to the mm10 genome using Bowtie (v1.0.1) (Langmead, Trapnell, Pop, & Salzberg, 2009), using options “-q --best --strata -m 1 --chunkmbs 1024”. Genome-wide TF binding events were called in each condition using MultiGPS (v0.74) (Mahony et al., 2014). EdgeR (v3.22.5) was used within the MultiGPS framework to test whether genomic sites were differentially bound by TFs (Robinson et al., 2010). Specifically, edgeR uses a negative binomial generalized linear model (GLM) to test whether ChIP-seq reads in one condition are significantly greater than in alternative condition. A genomic site was defined as shared by TFs if significant binding events were called in both TF ChIP-seq experiments (q-value < 0.001), and the TFs did not display differential read enrichment at that site as estimated by edgeR (q-value > 0.01). A binding event was defined as “TF1>TF2” if a MultiGPS peak was called in TF1 ChIP-seq (q-value < 0.001), TF1 exhibited a greater log fold-change with respect to the input ChIP-seq than TF2, and TF1 and TF2 were significantly differentially bound as defined by edgeR (q-value < 0.01). A similar strategy was applied to perform multi-way ChIP-seq comparisons. For example, when comparing HOXC6, HOXC9, and HOXC10 ChIP-seq experiments, “shared” binding sites were defined if significant binding events were detected in all three ChIP-seq datasets, and no two TFs were differentially bound with respect to each other. Sites were defined as “TF1> TF2, TF3” if a significant binding event was called for TF1, and a significantly greater number of reads were detected by EdgeR in TF1 ChIP-seq when compared to both TF2 and TF3. Finally, “TF1, TF2 > TF3” events were defined if significant binding events were called in both TF1 and TF2, no differential binding was detected between TF1 and TF2, and both TF1 and TF2 has a significantly greater number of reads than TF2. Only binding categories containing at least 500 binding events were retained.

### RNA-seq Data Processing

Fastq files obtained from RNA-sequencing were aligned to the genome using the splice-aware STAR (Spliced Transcripts Alignment to a Reference) aligner (v2.7.0c) (Dobin & Gingeras, 2016). Mapped reads were assigned to NCBI RefSeq annotated mm10 genes using the featureCount function in Rsubread (v1.30.9) (Liao, Smyth, & Shi, 2019). RefSeq genes with matching Entrez IDs were merged into a single gene by Rsubread. Following read summarization, read counts were normalized using the ‘rlog’ or regularized log transformation in DESeq2 (v1.20.0) (Love, Huber, & Anders, 2014). Transformed read counts were used as input features into the dimensionality reduction techniques PCA and MDS. The log_2_ fold change (LFC) in gene expression levels between iHox neurons vs. Day 2 progenitors or control neurons (no Dox treatment) was estimated using DESeq2. A q-value < 0.01 and LFC > 2 was used to define differentially expressed genes between Day 2 progenitors, control neurons, iHoxc6, iHoxc8, iHoxc9, and iHoxc10 neurons. We filtered out genes that were not expressed in any iHox neuron, retaining 19,019 genes.

### ATAC-seq Data Processing

Paired-end ATAC-seq reads were mapped to the mouse mm10 genome using Bowtie2 (v2.2.2) (Langmead et al., 2009). Genome-wide ATAC-seq derived accessible domains were defined using DomainFinder in the SeqCode project (https://github.com/seqcode/seqcode-core/blob/master/src/org/seqcode/projects/seed/DomainFinder.java).

### ChIP-seq and ATAC-seq Data Visualization

Heatmaps were used to plot the ChIP-seq and ATAC-seq reads at multiple categories of genomic sites in iHox neurons. Heatmaps were made using the MetaMaker program in SeqCode. (https://github.com/seqcode/seqcode-core/blob/master/src/org/seqcode/viz/metaprofile/MetaMaker.java). Raw reads from the ChIP-seq data were extended by 100 bps and read counts were binned into 100 bp bins. Binned reads were plotted over 1000 base pair windows centered on MultiGPS binding events. Color thresholds used to produce heatmaps were determined from binding data using the MetaPlotLineMaxEstimator program in SeqCode. Specifically, all 100 bp bins were ordered by the number of reads overlapping each bin. The maximum value for the heatmap color scale was set to the number of read counts at the 85^th^ percentile bin. The minimum value for all heatmaps was set to 5 reads. Finally, all binding events represented were ordered based on their genomic co-ordinates.

### ATAC-seq Composite Plots

For each ATAC-seq experiment, reads mapping to mitochondrial DNA or unannotated regions were filtered out. All experiments were performed in replicates; composite read distributions were initially calculated independently for each replicate. Filtered bedfiles were used to calculate the total number of reads overlapping each position in a 2000 base pair window surrounding a binding event of interest. The number of reads at each position was summed over the set of genomic sites being analyzed. The reads were total tag normalized, and finally the resulting read counts were averaged over replicates. Read density (reads per million per site or reads per million per 1000 sites) was plotted using seaborn in Python.

### RNA-seq data visualization

The log_2_ Fold Change (LFC) of gene expression levels in iHox neurons versus Day 2 progenitors or control neurons (no Dox treatment) was estimated using DESeq2. The LFC values were plotted for previously established marker genes in iHoxc6 neurons vs. Day 2 progenitors, iHoxc8 neurons vs. Day 2 progenitors, iHoxc9 neurons vs. Day 2 progenitors, iHoxc10 neurons vs. Day 2 progenitors, and control neurons vs. Day 2 progenitors (Figure 1D). The LFC values were plotted for previously established marker genes in iHoxc6 vs. control neurons, iHoxc8 vs. control neurons, iHoxc9 vs. control neurons, and iHoxc10 vs. control neurons (Supp Figure 2A). Volcano-style plots were used to simultaneously plot the LFC and p-values of significantly differentially expressed marker genes in the iHox versus control neuron comparisons (Supp Figure 2B). Volcano plots were used to represent the overall differential expression landscape between iHox and control neurons (Supp Figure 1C).

### Gene Ontology (GO) Term Enrichment Analysis

GREAT (Genomic Regions Enrichment of Annotation Tool, v4.0.4) (McLean et al., 2010) was used to perform GO term enrichment analysis. GREAT first calculates the probability that a randomly selected cis-regulatory element will be associated with a given GO term. It then uses a binomial test parameterized by this probability to determine whether a predefined subset of ChIP-seq peaks is significantly associated with that GO term. The process is repeated for all GO terms, and a list of significant associations is returned (McLean et al., 2010). We used GREAT to perform GO term enrichment analysis for the HOXC6-only, HOXC9-only, HOXC6&C9, HOXC9&C10, and shared HOX ChIP-seq binding sites. The top 10 significant GO terms were plotted; ordered by their Bonferroni corrected p-values.

### Association between gene sets and binding events

In order to construct gene sets, pairwise comparisons of gene expression levels between iHoxc6, iHoxc9 and iHoxc10 neurons vs. Day 2 progenitors were performed using DESeq2. Additionally, pairwise comparisons were also performed between the iHox neurons: iHoxc6 vs. iHoxc9, iHoxc6 vs. iHoxc10 and iHoxc9 vs. iHoxc10. Genes that were up-regulated in all iHox vs. Day 2 progenitor comparisons (log_2_ fold change (LFC) > 2 and adjusted p-value < 0.01), as well as not differentially expressed between the iHox neurons (|LFC| > 2) were assigned to the gene set “shared-upregulated”. A similar logic was used to identify the “shared-downregulated” genes. Pairwise comparisons between iHox neurons were used to identify genes significantly up-regulated or downregulated (|LFC| > 2 and adjusted p-value < 0.01) in individual iHox neurons. Genes that were significantly up-regulated or downregulated in two out of three iHox neurons were assigned to gene sets using the same thresholds.

In order to test whether specific categories of ChIP-seq binding events were associated with differentially regulated gene sets, we used ChIP-Enrich (v.1.10.0). Specifically, each peak was assigned to the gene with the nearest TSS. ChIP-Enrich uses a logistic regression model to test whether gene set membership (controlled by mappable gene length) predicts whether a gene is associated with a ChIP-seq peak (Welch et al., 2014). The significance of the weight associated with gene set membership is estimated using the ‘Wald’ statistic. P-values obtained from the Wald statistic adjusted using the Benjamini-Hochberg multiple testing correction (Welch et al., 2014). For each category of binding events, adjusted p-values were plotted for each pre-defined gene set.

### DNA motif analysis

*De novo* motif discovery was performed using MEME-ChIP (v. 5.1.0) (Machanick & Bailey, 2011) with the following command-line settings: “-ccut 100 -dna -meme-p 4 -meme-mod anr -meme-minw 5 -meme-maxw 15 -meme-nmotifs 10 -dreme-e 0.05”. MEME-ChIP was also provided knowledge of the JASPAR CORE motif database (2016 release) (Mathelier et al., 2016) via the “-db” option for the purposes of TOMTOM motif similarity matching and CentriMo analysis. To find *de novo* discovered motifs that matched canonical cognate HOX TF binding motifs, we first extracted MEME or DREME discovered motifs that received significance scores of less than 1e-3 from those tools. We then used STAMP (v. 1.0) (Mahony & Benos, 2007) with settings “-cc PCC -align SWU” to match against the following motif consensus sequences: TAATDR, HHATAAA, TAAT, ATAAA, GTAAA, TAAAC, TAAAT. Motifs were retained as probable cognate HOX motifs if they matched one of the above consensus sequences with a STAMP E-value score of less than 1e-4. Further motif frequency analysis was performed for two motifs that were discovered by DREME in the c6>c9,c10 and c9>c6,c10 categories (Figure 3B). A motif scanning procedure using log likelihood scoring was used to find peaks that contained motif hits within 50bp of the peak positions. Motif hits were defined as sequences scoring above motif scanning thresholds set using a 0.1 FDR derived from 1 million sequences (100bp) randomly generated from a 2^nd^-order Markov model of the mouse genome. Over-representation of peaks containing motifs was assessed in comparison to 10,000 sequences (100bp) randomly sampled from the mouse genome. Finally, SeqUnwinder (v. 0.1.3) (Kakumanu et al., 2017) was used to find motifs that discriminate between various HOX binding site categories using the following command-line settings: “--threads 10 --makerandregs --makerandregs --win 150 --mink 4 --maxk 5 --r 10 --x 3 --a 400 --hillsthresh 0.1 --memesearchwin 16”.

### Bichrom Data Analysis

For each HOX TF, we first trained a hybrid convolutional and long short-term memory neural network (CNN-LSTM) to predict induced HOX binding using only DNA sequence information. We then applied Bichrom to the HOX TFs, which integrates DNA sequence and preexisting chromatin to predict induced TF binding. In particular, Bichrom takes the trained sequence-only network and incorporates additional chromatin data using a secondary chromatin sub-network (Srivastava et al., 2019). Day 2 progenitor ATAC-seq, H3K27ac, H3K4me3, H3K9me3, H3K27me3 and PolII were used to define the preexisting chromatin landscape. For each chromatin experiment, the tag counts at each genomic window were total tag normalized, averaged across replicates and binned into 50 base pair bins. The binned chromatin tracks were used as inputs into Bichrom’s chromatin sub-network. To account for computational variation in network performance, we repeated the training process on 9 distinct training sets, each corresponding to a separate held-out chromosome. We used the area under the precision-recall curve (auPRC) to measure the genome-wide predictive performance of the sequence-only neural network and Bichrom on the 7 HOX TFs. The hyper-parameters for all networks were selected using a hyper-parameter grid search, and the networks were built using Keras (https://github.com/seqcode/iTF).

---

**Supp Figure 1:**
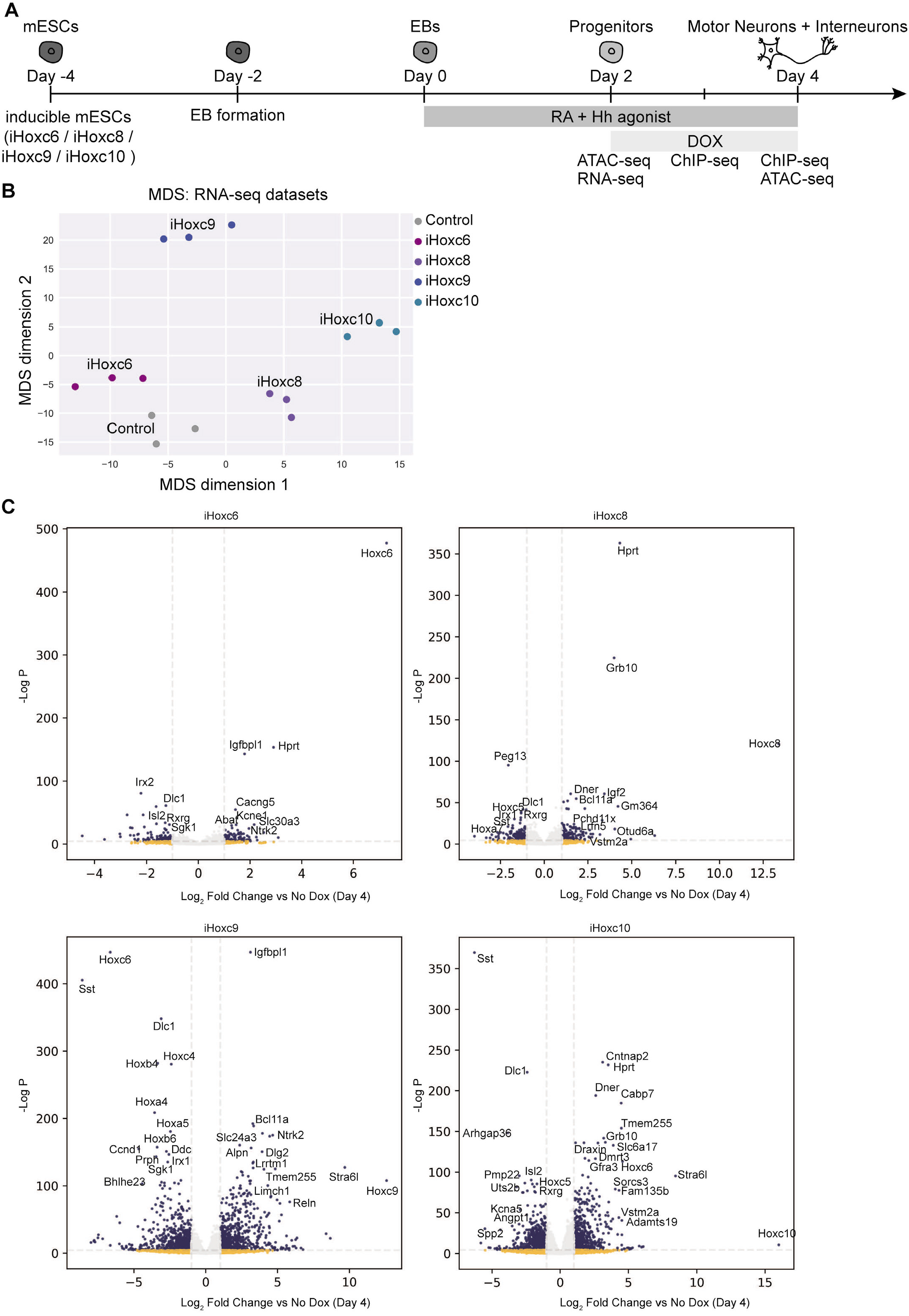
HOX TFs induce distinct gene expression profiles. (A) Experimental scheme. (mESCs: mouse embryonic stem cells. EBs: embryoid bodies) (B) Multidimensional scaling (MDS) of the RNA-seq datasets (Day 4) reveals similarities in the gene expression profiles induced by HOX TFs. (C) Volcano plots showing changes in gene expression in iHoxc6, iHoxc8, iHoxc9 and iHoxc10 relative to No Dox control neurons (Day 4).

**Supp Figure 2:**
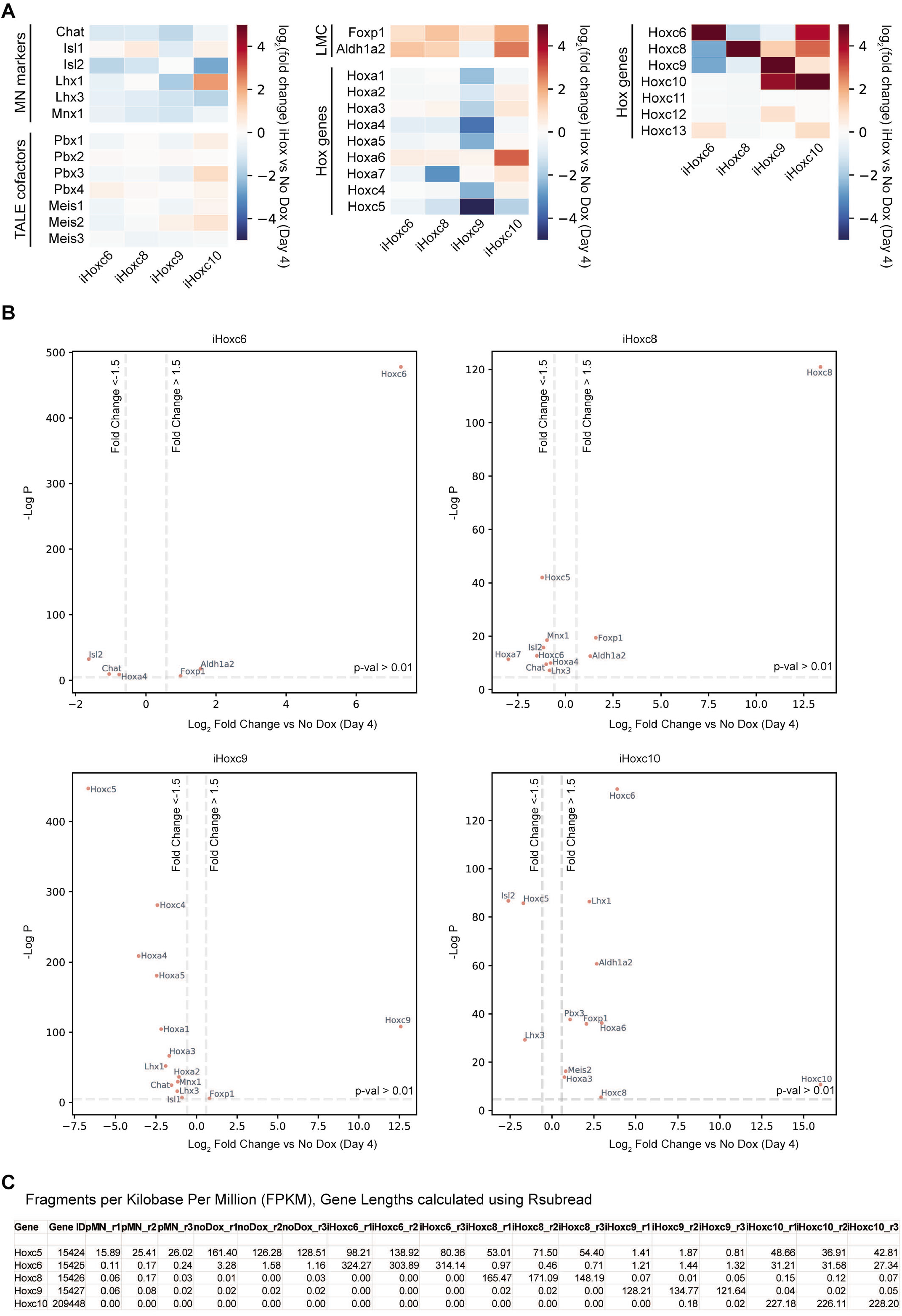
HOX TFs induce the expected markers during *in vitro* spinal cord differentiation. (A) RNA–seq heatmap showing the expression of representative genes in iHoxc6, iHoxc8, iHoxc9 and iHoxc10 relative to No Dox control neurons (Day 4). (B) Plots showing significantly differentially expressed representative genes from A. (C) Normalized (FPKM) read counts for *Hox* genes in the indicated conditions (in 3 biological replicates) (pMN: progenitors (Day 2)).

**Supp Figure 3:**
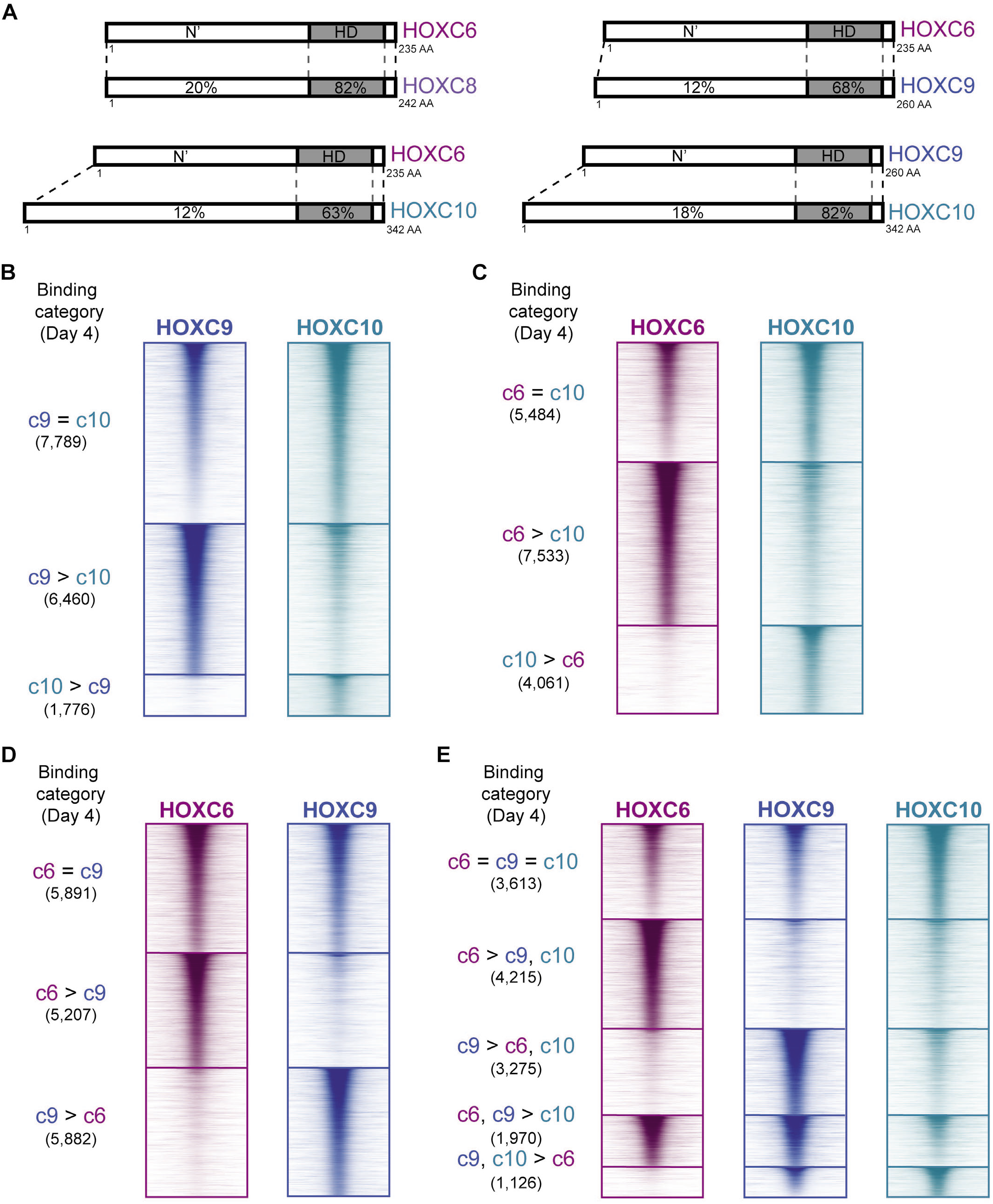
HOXC6, HOXC9, and HOXC10 TFs have different genome-wide binding profiles at Day 4. (A) Schematic indicating the percentage of conserved amino acids in the homeodomain (HD) and total protein between the indicated HOX TFs. (B-D) ChIP-seq heatmap showing binding comparisons in differentiating neurons, at Day 4. Sites bound by both indicated HOX TFs noted as “=” sites. Preferentially bound sites by HOXC6, HOXC9 or HOXC10 noted as “c6 >”, “c9 >”, and “c10 >”, respectively. (E) ChIP-seq heatmap showing binding comparisons of HOXC6, HOXC9, and HOXC10 in differentiating neurons, at Day 4. Sites bound by all three HOX TFs noted as “c6 = c9 = c10” sites. Preferentially bound sites by HOXC6, HOXC9, HOXC6 and HOXC9 or HOXC9 and HOXC10 noted as “c6 > c9, c10”, “c9 > c6, c10”, “c6, c9 > c10”, and “c9, c10 > c6”, respectively.

**Supp Figure 4:**
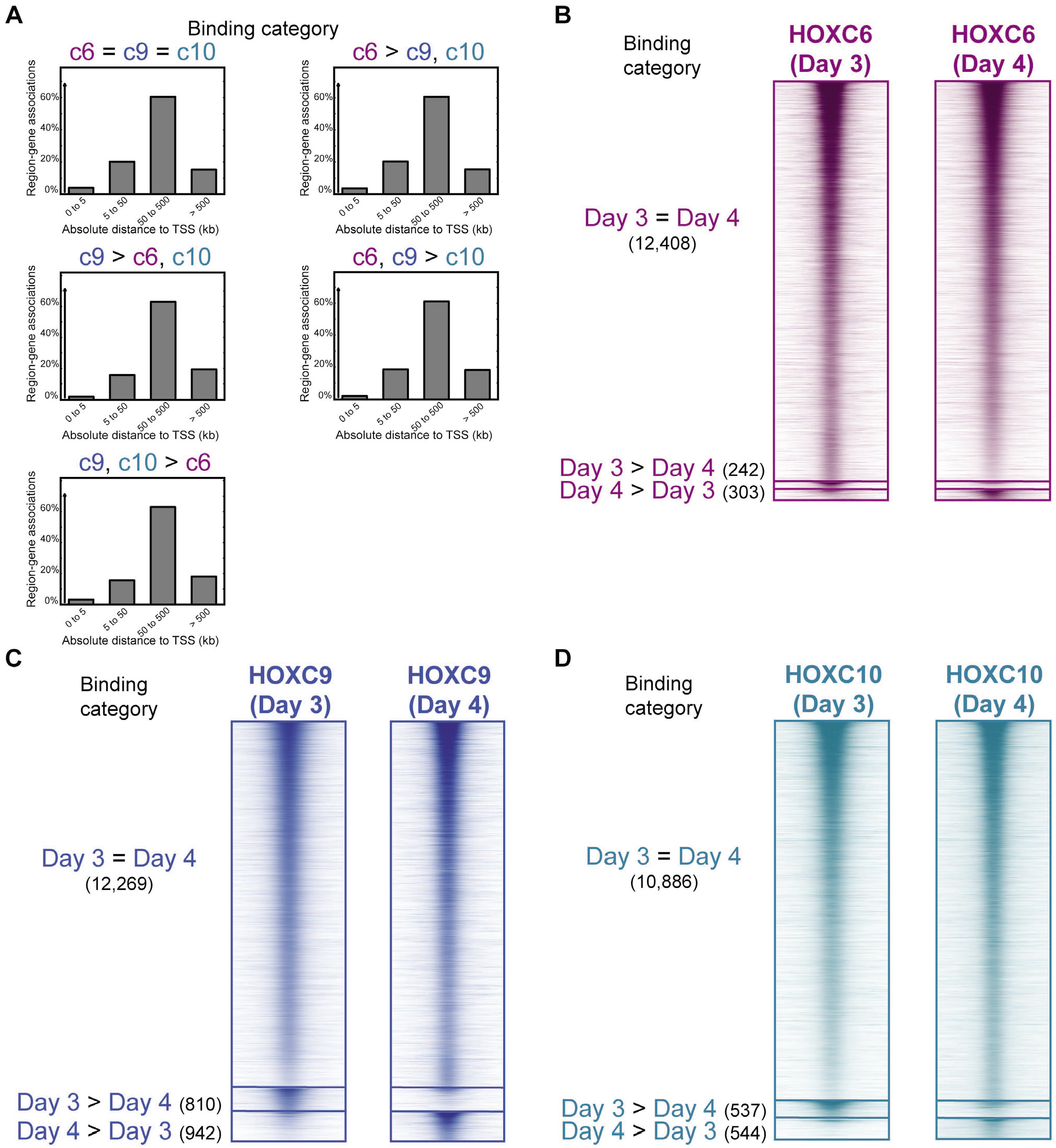
HOXC6, HOXC9, and HOXC10 TFs have largely the same binding patterns at Day 3 vs Day 4. (A) Region-gene association graphs at the indicated HOX binding categories from Figure 2E. (B-D) ChIP-seq heatmaps showing binding comparisons of the indicated HOX TFs at Day 3 versus Day 4. Sites equally bound at both time points noted as “Day 3 = Day 4” sites. Differentially bound sites noted as “Day 3 > Day 4” and “Day 4 > Day 3”.

**Supp Figure 5:**
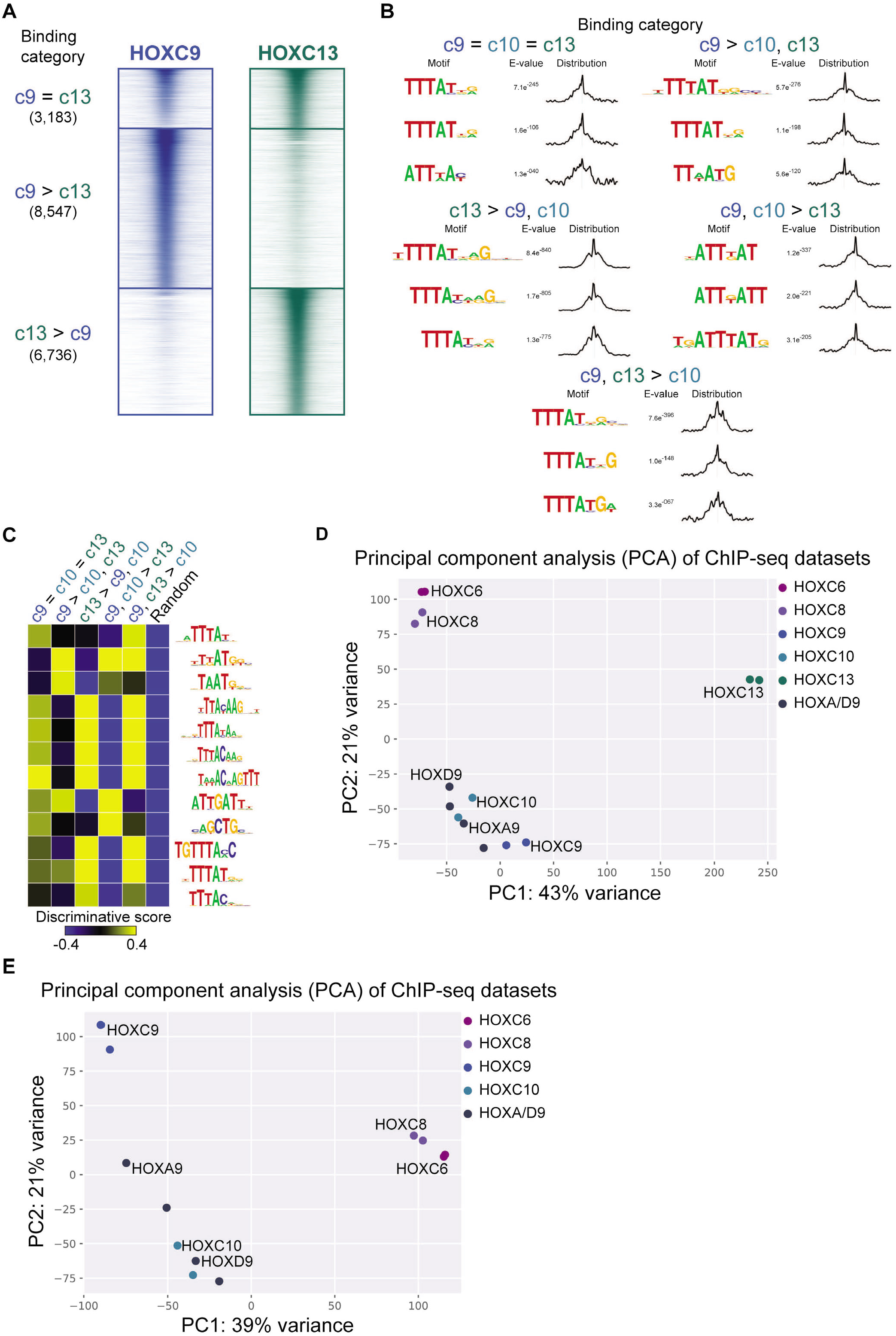
HOXC9 and HOXC13 genome-wide binding profiles differ from other posterior HOX TFs. (A) ChIP-seq heatmap showing binding comparisons of HOXC9 and HOXC13 in differentiating neurons, at Day 3. Sites bound by both indicated HOX TFs noted as “c9 = c13” sites. Preferentially bound sites by HOXC9 or HOXC13 noted as “c9 > c13” or “c13 > c9“. (B) Selected top enriched motifs discovered via MEME-ChIP at the indicated HOX binding categories from Figure 7A. Distributions to the left of each motifs show the distribution of each motif occurrence with respect to the midpoint of each peak (500bp windows). (C) SeqUnwinder analysis characterizing motifs that are discriminative between the various classes of HOXC9, HOXC10, and HOXC13 binding sites. (D-E) Principal Component Analysis (PCA) of the ChIP-seq datasets reveals similarities in the binding patterns of HOX TFs.

**Supp Figure 6:**
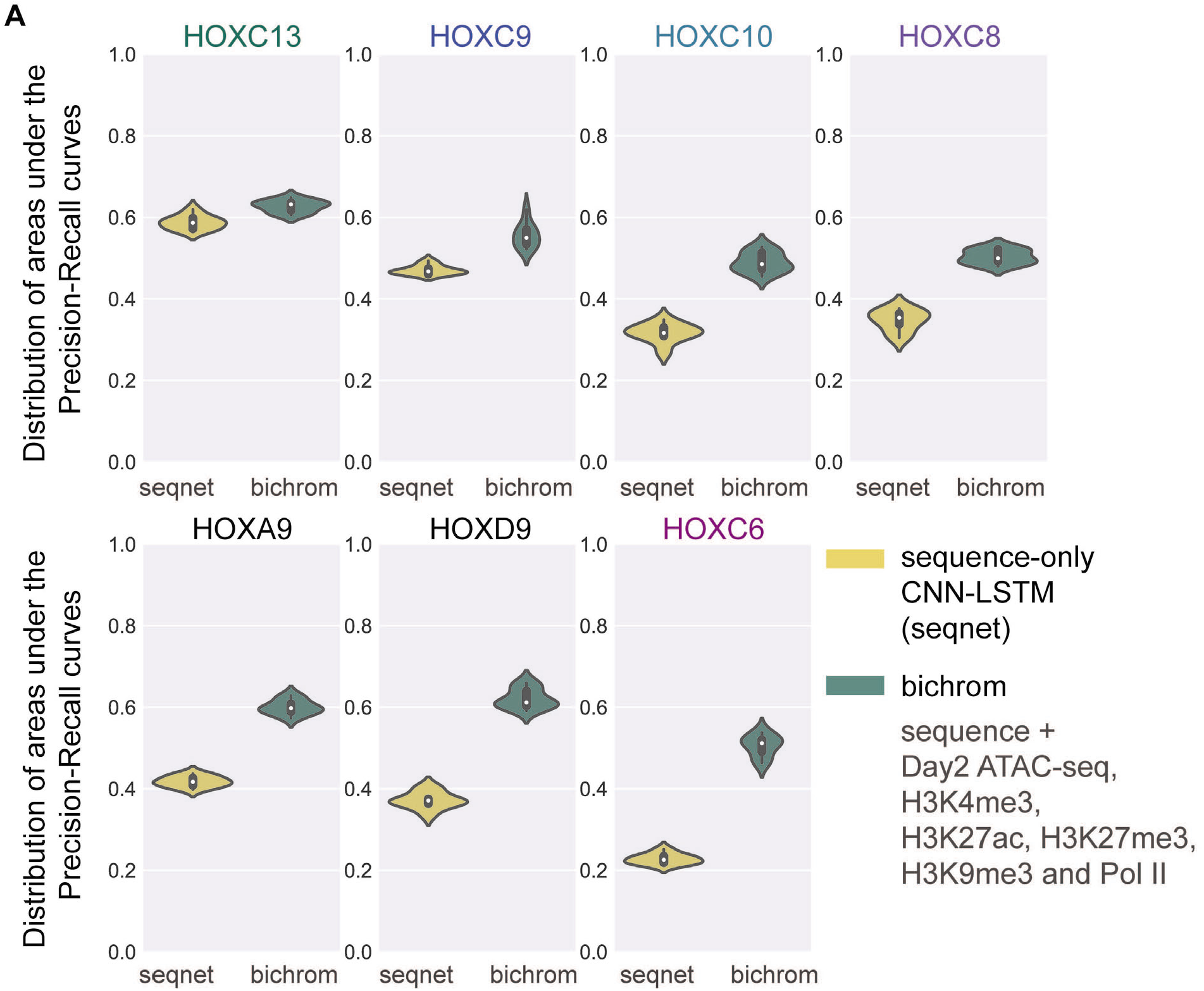
Neural network-based analysis quantifies HOX TFs’ relative dependence on prior chromatin states. (A) The distribution of areas under the precision-recall curve (auPRCs) for 9 iterations of a CNN-LSTM sequence-only network (seqnet, yellow) compared to a bimodal network that incorporates both sequence and Day 2 (progenitors) chromatin data to predict genome-wide induced HOX TF binding (Bichrom, green). The chromatin data used includes Day 2 ATAC-seq and ChIP-seq for H3K4me3, H3K27ac, H3K27me3, H3K9me3 and Pol II. For each TF, both seqnet and Bichrom are trained using 9 distinct training sets, each corresponding to an independent held out test-set. The auPRC distributions are represented using violin plots.

**Supp Figure 7:**
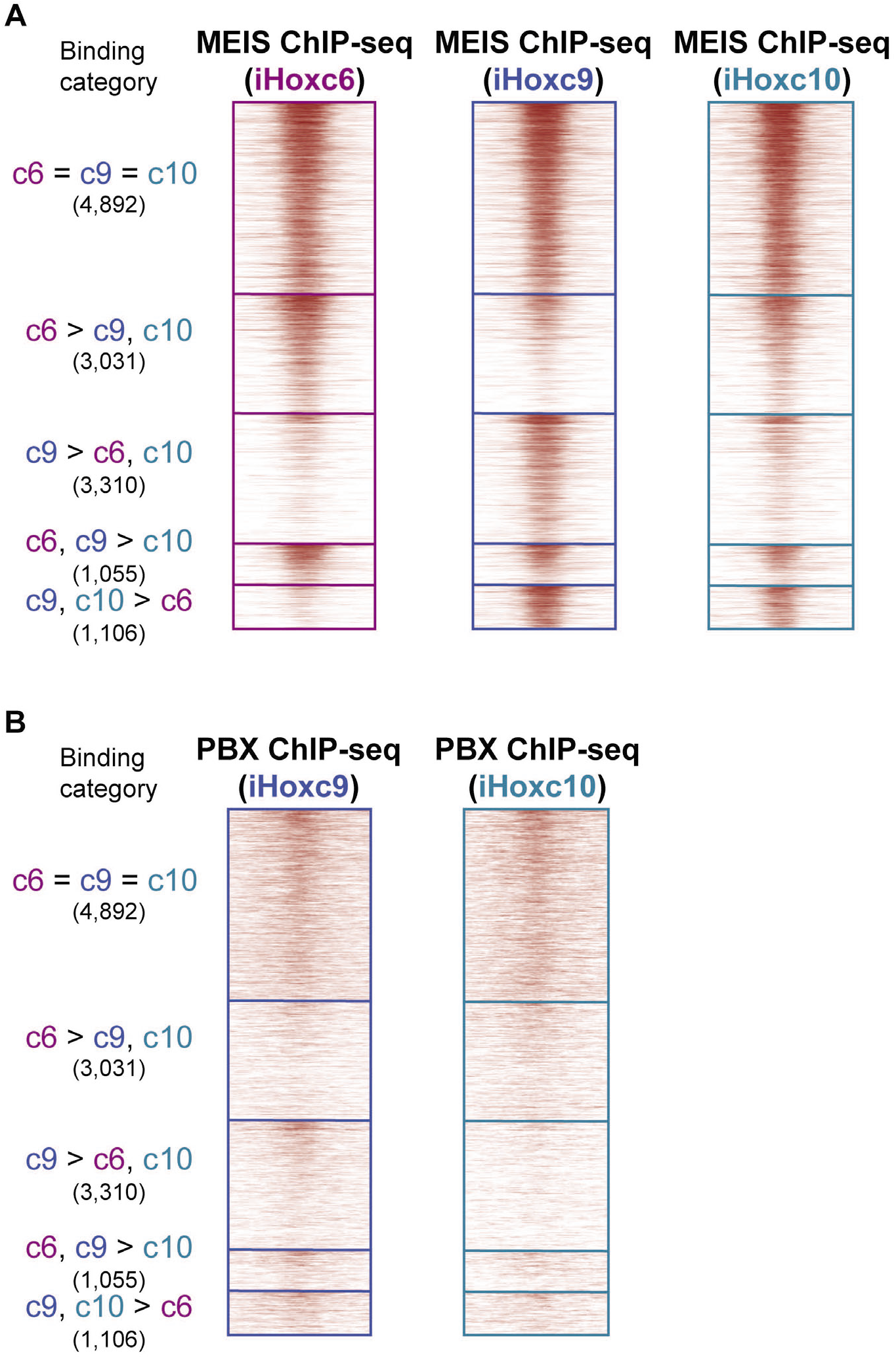
Overlap of TALE cofactor binding with HOXC6, HOXC9, and HOXC10. (A-B) ChIP-seq of MEIS and PBX performed in the indicated cell line reveals co-binding with the respective HOX TF at the indicated HOX binding categories from Figure 2E.

## Notes

### Competing Interest Statement

The authors have declared no competing interest.

### Summary of Updates

We have added an analysis that supports further the original results. Bichrom is a neural network-based method that integrates DNA sequence and prior chromatin information to explain the observed genomic binding patterns of an induced TF. We train Bichrom to predict the binding patterns of each HOX TF using DNA sequence features and prior chromatin features. Following training, we compared the performance of the sequence-only network to that of Bichrom on held-out test sets. We find that HOXC9 and HOXC13 networks display minor improvements in the predictive performance when trained with or without neuronal progenitor chromatin data. Consistent with our previous results, none of the various incorporated prior chromatin tracks predict future binding for those posterior TFs. HOXC10 and HOXC8 predictions benefit from included prior chromatin data, but HOXC6 and the other HOX9 paralogs (HOXA9 and HOXD9) display substantial gains in predictive performance when training includes preceding chromatin tracks. These results support the hypothesis that even HOX TFs from the same group rely on previous chromatin states to different degrees for their genomic binding.

